# Molecular Structure and Enzymatic Mechanism of the Human Collagen Hydroxylysine Galactosyltransferase GLT25D1/COLGALT1

**DOI:** 10.1101/2024.06.21.600124

**Authors:** Matteo De Marco, Sristi Raj Rai, Luigi Scietti, Daiana Mattoteia, Stefano Liberi, Alberta Pinnola, Federico Forneris

## Abstract

During biosynthesis, collagen lysine residues undergo extensive post-translational modifications essential for the stability and functions of collagen supramolecular assemblies. In the endoplasmic reticulum, two distinct metal ion dependent enzyme families (i.e., multifunctional lysyl hydroxylases-glucosyltransferases LH/PLODs and galactosyltransferases GLT25D/COLGALT) alternatively operate on collagen lysine side chains ultimately generating the highly conserved α-(1,2)-glucosyl-β-(1,O)-galactosyl-5-hydroxylysine pattern. Malfunctions in the collagen lysine post-translational modification machinery is linked to multiple developmental pathologies as well as extracellular matrix alterations causing enhanced cellular proliferation and invasiveness of several solid tumors, prompting for an in-depth characterization of LH/PLOD and GLT25D/COLGALT enzyme families. Here, we present an integrative molecular study of GLT25D1/COLGALT, highlighting an elongated head-to-head homodimeric assembly characterized by an N-terminal segment of each monomer wrapping around its dimerization partner. Each monomer encompasses two Rossman fold-type domains (GT1 and GT2) separated by an extended linker. Both domains were found capable of binding Mn^2+^ cofactors and UDP-α-galactose donor substrates, resulting in four candidate catalytic sites per dimer. Site-directed mutagenesis and biochemical studies identify the C-terminal GT2 domain as the functional GLT25D1/COLGALT1 catalytic site, highlighting an unprecedented Glu-Asp-Asp motif critical for metal ion binding, and suggesting structural roles for the N-terminal GT1 essential for correct quaternary structure assembly. Conversely, dimerization was not a requirement for GLT25D1/COLGALT1 enzymatic activity *in vitro*, suggesting that the elongated enzyme homodimer assembly, resembling that of LH/PLOD binding partners, could represent a functional hallmark for correct recognition and successful processing of collagen lysine residues.

## INTRODUCTION

Collagens are highly conserved proteins, from invertebrates to higher vertebrates. Their amino acid composition, post-translational modification patterns, and supramolecular assembly confer a versatile spectrum of physical and mechanical properties, resulting in the most abundant protein family in the animal kingdom constituting essential components of several animal tissues including bones, cartilage, tendons, and dermis^1–3^. During biosynthesis, collagens undergo extensive post-translational modifications to fold into right-handed triple helices. The repetitive unit consists of the triplet Gly-X-Y, where positions X and Y are frequently occupied by proline and lysine residues^4–7^. The role of proline hydroxylation in the global process of collagen biosynthesis is the most studied and understood to date, essential to the winding process of the collagen right-handed triple helix^8–10^. Contrary to proline residues, the role of lysine residues is mainly linked to the cross-linking processes that occur between collagen molecules once secreted into the extracellular space^11,12^.

Lysine residues in collagens and collagen-like protein segments are modified sequentially by specific biosynthetic enzymes which catalyze the formation of a simple but very conserved post-translational modification pattern: the α-(1,2)-glucosyl-β-(1,O)-galactosyl-5-hydroxylysine (Glc-Gal-Hyl)^12–16^. The path to the formation of Glc-Gal-Hyl starts with the formation of 5-hydroxylysine (Hyl) operated by collagen lysyl hydroxylases (LH, also known as procollagen, lysyl 2-oxoglutarate dioxygenases (PLOD)). The LH/PLOD enzyme family is characterized in humans by three different isoforms (LH1/PLOD1, LH2/PLOD2 and LH3/PLOD3)^11,17,18^. The LH reaction depends on the presence of tightly bound Fe^2+^ in the C-terminal domain of LH/PLODs, which is essential for the electron transfer, and on ascorbate, that preserves the redox state of the metal ion^19–21^. Hyl residues are then glycosylated by Mn^2+^-dependent galactosyltransferase enzymes^6,22,23^, through an inverting reaction that adds the galactose moiety of the uridine diphosphate-α-galactose (UDP-α-Gal) donor substrate to the Hyl hydroxyl group, generating β-(1,O)-5-hydroxylysine (Gal-T activity). Gal-Hyl is further glycosylated by the multifunctional LH/PLOD enzymes through a retaining reaction, that requires Mn^2+^ and uridine diphosphate-α-glucose (UDP-α-Glc) donor substrate (Glc-T activity), ultimately generating the Glc-Gal-Hyl. Initially, the multifunctional LH/PLOD enzyme(s) were thought to be responsible for the entire Lys–to–Glc-Gal-Hyl biosynthetic pathway^24,25^. The subsequent identification of a specific family of collagen galactosyltranfserases (named GLT25D or COLGALT)^26,27^, underpinned their essential roles in collagen biosynthesis *in vivo* ^23,28–32^. Additional biochemical studies on human LH/PLOD and GLT25D/COLGALT enzymes^27,33^ ultimately ruled out the single enzyme hypothesis. The resulting picture shows how the concerted action of two enzyme systems (i.e., LH/PLOD and GLT25D/COLGALT) alternatively acting on the same substrate and possibly directly interacting with each other, lead to complete Glc-Gal-Hyl collagen post-translational modifications (Fig. 1a).

**Figure 1.**
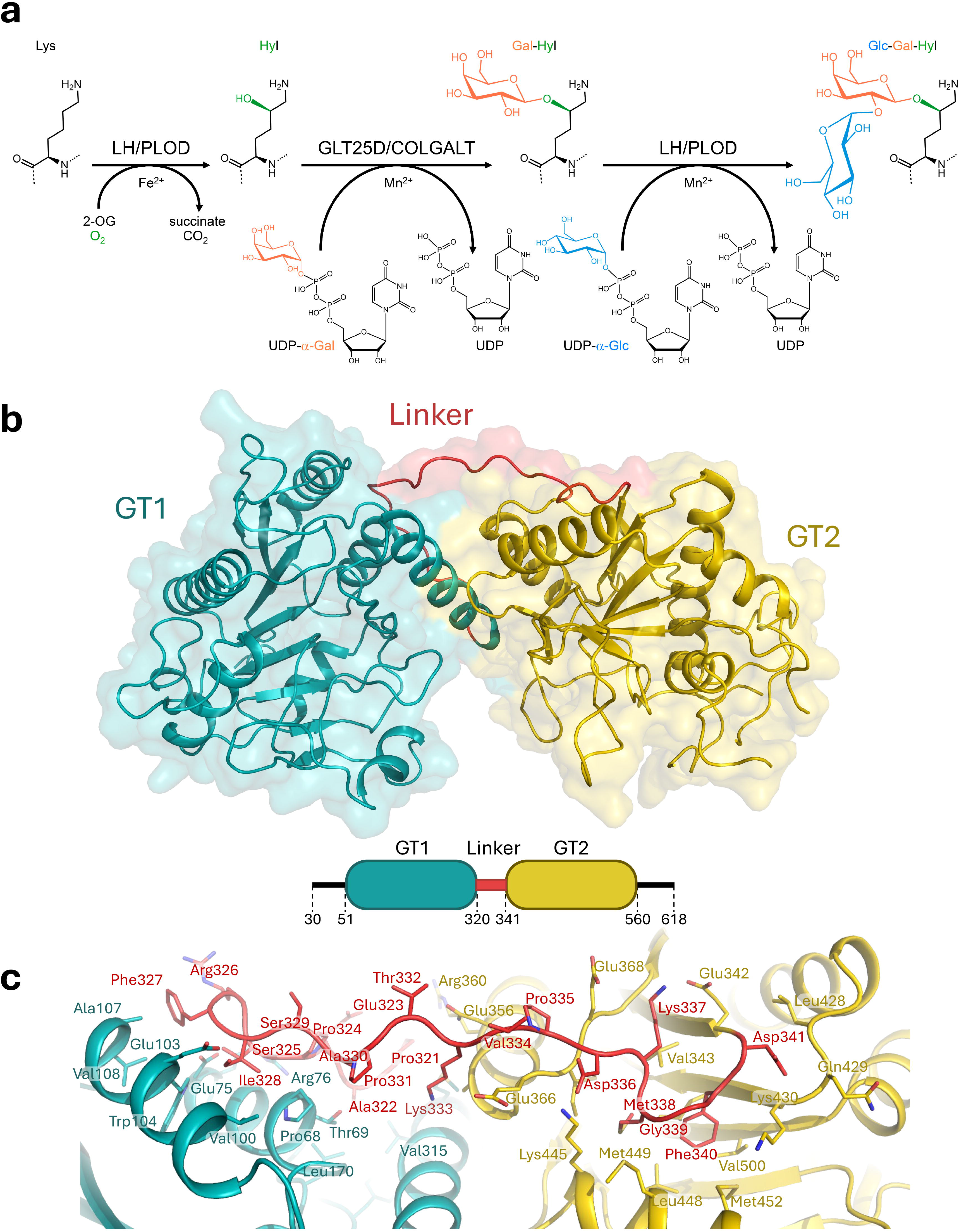
Biochemical and structural features of human GLT25D1/COLGALT1. (a) Reaction scheme showing the collagen Lys–to–Glc-Gal-Hyl conversion post-translational modification process mediated by the alternate action of LH/PLOD and GLT25D1/COLGALT1, with color highlight of donor substrates and associated products; (b) Cartoon representation of the crystal structure of human GLT25D1/COLGALT1, showing the domain architecture of the enzyme with GT1 and GT2 domains connected via a rigid linker element, anchored to both domains via hydrophobic anchorages shown in details in panel (c).

While multifunctional lysyl hydroxylase and glucosyltransferase LH/PLOD enzymes have been subject to extensive molecular, structural and characterizations^33–36^, much less is known regarding the GLT25D/COLGALT enzymes responsible for the central galactosyltransferase reaction. Human GLT25D/COLGALT are soluble proteins anchored to the endoplasmic reticulum through a C-terminal Arg-Asp-Glu-Leu (RDEL) sequence, and have been found to co-localize with LH/PLOD enzymes^26,27^. Three human GLT25D/COLGALT isoforms have been identified: GLT25D1/COLGALT1, constitutively present in all human tissues, GLT25D2/COLGALT2, expressed at the level of nervous tissue, and GLT25D3/COLGALT3, found in all secretory tissues such as salivary glands, pancreas, placenta, as well as in the nervous system. However, only the first two isoforms are active, whereas isoform 3 (also called CERCAM) is inactive *in vitro* and potentially not related at all to collagen lysine modification^37–39^.

Of considerable relevance is the involvement of the *colgalt* genes and their protein products GLT25D/COLGALT enzymes in numerous pathological conditions, including cerebral small vessel disease^40^, liver pathologies^41^, ostheoarthritis^32,42,43^, musculoskeletal defects^28^, and cancer^44,45^. Liver fibrosis is a chronic pathological condition that sees an accumulation of proteins at the level of the extracellular matrix, whose evolution often leads affected subjects to develop cirrhosis and hepatocellular carcinoma. Recent work has observed high levels of GLT25D1/COLGALT1 in the plasma of cirrhosis patients compared to unaffected patients, and one of the proposed hypotheses is the stimulatory role of the enzyme against hepatic stellate cells ^46^.

Efforts to decipher the structure-function relationships underlying GLT25D/COLGALT activity on collagen molecules yielded interesting insights^37^, but highlighted the presence of multiple Asp-x-Asp (DxD) motifs along the polypeptide sequence potentially relevant for enzymatic activity, thus demanding more in-depth structure-based investigations. Here, we merged structural biology methods with biochemistry and biophysics to reveal the quaternary assembly of GLT25D1/COLGALT1. The X-ray crystal structures of full-length human GLT25D1/COLGALT1 in its native state as well as in complex with the donor substrate UDP-α-Gal were determined using experimental phasing. Small-angle X-ray scattering, negative stain electron microscopy and mass photometry jointly revealed an elongated quaternary structure reminiscent of that of LH/PLOD enzymes, characterized by a head-to-head dimer with each monomer composed of two similar domains with Rossman fold topology, both populated by the Mn^2+^ cofactor and the donor substrate. Structure-guided site-directed mutagenesis ultimately identified the C-terminal domain as the functional site for enzyme catalysis, while the N-terminal domain retains substrates and cofactors exclusively for folding stability purposes. Collectively, our results provide insights on the molecular architecture and functions of a key human enzyme of collagen biosynthesis.

## MATERIALS AND METHODS

### Chemicals

All chemicals were purchased from Sigma-Aldrich unless otherwise specified.

### DNA constructs

The sequence encoding for human GLT25D1/COLGALT1, deprived of the N-terminal signal peptide and the C-terminal RDEL retention sequence (UniProt Q8NBJ5, residues 30 to 618), was synthesised by Genewiz and sub-cloned into a pCR8 cloning vector with in-frame 5’-BamHI and 3’-NotI restriction sites. GLT25D1/COLGALT1 mutants were generated using the Phusion Site Directed Mutagenesis Kit (Thermo Fisher Scientific). The entire plasmid was amplified using the primers listed in Supplementary Table 1. The linear mutagenized plasmids were then phosphorylated using T4 polynucleotide kinase (Invitrogen) prior to ligation. All plasmids were checked by Sanger sequencing (Microsynth) prior to cloning into a modified pET28b-SUMO vector (Novagen). The recombinant plasmids include an N-terminal 8xHis-tag followed by a small ubiquitin-like modifier (SUMO) tag preceding the in-frame 5’-BamHI restriction site, as well as an in-frame stop codon after the 3’-NotI restriction site.

### Production of recombinant GLT25D1/COLGALT1

Chemically competent *E. coli* BL21 cells (Invitrogen) were transformed with the pET28b-SUMO-GLT25D1 recombinant plasmid using heat shock. A single colony was inoculated into Lysogeny broth (LB) medium supplemented with 100 μg/ml of kanamycin and grown overnight at 37 °C in a shaking incubator (New Brunswick). This pre-culture was then inoculated at a 1:50 ratio in 1 L of ZYP5052 autoinducing medium^47^ in a 5 L Erlenmeyer flask. The culture was kept at 37 °C for 3 hours at 180 r.p.m. shaking speed. The temperature was then lowered to 17 °C, and further incubated overnight. Bacterial cells were harvested by centrifugation at 5000 *g*, resuspended at 1:10 *w/v* ratio in a lysis buffer composed of 25 mM 4-(2-hydroxyethyl)-1-piperazineethanesulfonic acid (HEPES)/NaOH, 500 mM NaCl, 10 μM leupeptin, 10 μM pepstatin, 0.3 mg/ml chicken egg white lysozyme, 500 μM MnSO_4_, pH 8.0. The suspension was subject to sonication (16 cycles, 9 s on, 6 s off pulses). Cell debris was removed by centrifuging the cell lysate at 60,000 *g* for 45 min, at 4 °C in an Avanti J26 super centrifuge (Beckman Coulter). The supernatant containing soluble His-SUMO-GLT25D1/COLGALT1 protein was filtered through a MiniSart GF 0.8 μm filter (Sartorius), loaded onto a 5 mL NiSepharose Excel column (Cytiva) and eluted using a stepwise gradient of an elution buffer composed of 25 mM HEPES/NaOH, 500 mM NaCl, 500 mM imidazole, pH 8.0. The fractions of the eluate containing His-SUMO-GLT25D1/COLGALT1, as assessed by SDS-PAGE analysis, were pooled and dialyzed overnight against 2 L of 25 mM HEPES/NaOH, 500 mM NaCl, pH 8.0. The N-terminal 8xHis-SUMO tag was simultaneously cleaved by incubating the protein with 1 mg/mL His-tagged SUMO protease (1:300 *v/v*). Affinity-based removal of SUMO protease and 8xHis-SUMO-tag was achieved by passing the purified sample through a 5 mL Ni Sepharose Excel column (Cytiva) and collecting the flow-through fraction. The resulting sample was subject to buffer exchange in 30 kDa MWCO Vivaspin Turbo centrifugal filters (Sartorius) against 25 mM HEPES/NaOH, 100 mM NaCl, pH 8.0 and loaded into a HiScreen Capto Q column (Cytiva) pre-equilibrated with the same buffer. A linear NaCl gradient was then applied, resulting in GLT25D1/COLGALT1 elution with approximately 250 mM NaCl. The sample was concentrated using 30 kDa MWCO Vivaspin Turbo centrifugal filters (Sartorius) and subject to final polishing into a Superdex 200 Increase 10/300 GL (Cytiva), equilibrated with 25 mM HEPES/NaOH, 100 mM NaCl, pH 8.0. The final sample quality was assessed by reducing and non-reducing SDS-PAGE analysis. GLT25D1/COLGALT1-containing fractions were pooled, concentrated to 4 mg/mL, and stored at -80 °C prior to usage.

### Protein Crystallization

GLT25D1/COLGALT1 spherulites were initially found in various conditions of nanoliter-dispensed droplets (0.1 μL protein at 4 mg/mL+ 0.1 μL reservoirs, dispensed using a Gryphon crystallization robot (Art Robbins) using the Morpheus (Molecular Dimensions) crystallization screen in sitting-drop vapor diffusion drop plates (SwissSci). After extensive optimization, diffraction-quality crystals were obtained by manually mixing 0.5 μL of protein concentrated at 3.5 mg/mL and 0.5 μL of a reservoir solution composed of 8% poly-ethylene-glycol (PEG) MW 4,000, 100 mM 2-(N-morpholino)ethanesulfonic acid (MES)/NaOH, 20% glycerol, pH 6.5. Co-crystallization experiments were performed by setting up the same conditions and supplementing the protein solution with 500 μM MnCl_2_ and 1 mM UDP-α-Gal. Crystals were harvested using mounted Litholoops (Molecular Dimensions), flash-cooled and stored in liquid nitrogen prior to data acquisition. Heavy atom derivatives were prepared by soaking the GLT25D1/COLGALT1 crystals in mother liquor conditions containing 1 mM K_2_HgBr_4_. Crystals were incubated with the heavy atom solution for at least 5 hours at 4 °C prior to cryoprotection, harvesting and flash-cooling in liquid nitrogen.

### X-ray data collection, structure determination and refinement

X-ray diffraction data from single crystals were collected at 100 K at various beamlines of the European Synchrotron Radiation Facility (ESRF) in Grenoble, France and the Swiss Light Source (SLS) in Villigen, Switzerland. Experimental phasing was achieved by collecting X-ray diffraction data at the mercury inflection point at the ID23-1 beamline of the ESRF synchrotron equipped with an Eiger 2 16M (Dectris) detector. Data were indexed and integrated with *XDS*^48^, followed by scaling and merging using *AIMLESS*^49^. Data collection statistics are given in Supplementary Table 2. Single wavelength anomalous dispersion (SAD) phasing was performed with the *SHELXC/D/E* pipeline^50^ and *HKL2MAP*^51^. A total of 13 heavy atom sites were identified at 4.0 Å resolution in *SHELXD*. Phasing with *SHELXE* resulted in non-ambiguous identification of the map handedness immediately after the initial rounds of density modification with phase extension to 2.80 Å, with 62% solvent content. The density modified experimental electron density map was used for tracing the initial model by combining multiple rounds of *ARP/wARP*^52^ and *BUCCANEER*^53^. The model was completed and the structure was refined by alternating steps of manual building in *COOT*^54^ and automated refinement with *phenix.refine*^55^ and *REFMAC5*^56^. Crystal structures of GLT25D1/COLGALT1 in complex with donor substrates were determined using molecular replacement in *PHASER*^57^, using the experimental structure of the enzyme as search model. Validation was carried out using *MolProbity*^58^ and the validation tools available on the Protein Data Bank server^59^. Final refinement statistics are summarized in Supplementary Table 3. Structural figures were prepared with *PyMol* (http://www.pymol.org). Superpositions were performed using the “*super*” command in *PyMol*. All-atom root mean square deviation (RMSD) values were computed accordingly.

### Mass Photometry

Mass photometry measurements were carried out on a Refeyn Two Mass Photometer (Refeyn) using 24 x 50 mm^2^ glass coverslips (Refeyn). Initial 200 nM stocks of purified human GLT25D1/COLGALT1 wild-type and mutants were diluted 1:10 using 25 mM HEPES/NaOH, 100mM NaCl, pH 8.0, and analyzed under a field of view of 4 μm x 11 μm at a frame rate of 500 Hz. The data were collected using *AcquireMP* (Refeyn), then processed and plotted using *DiscoverMP* (Refeyn). Results of the data analysis are summarized in Supplementary Table 4.

### Size Exclusion Chromatography coupled with Multi-Angle Light Scattering (SEC-MALS)

30 μl of 4 mg/mL recombinant GLT25D1/COLGALT1 were injected into a Protein KW-802.5 analytical size-exclusion column (Shodex) and separated with a flow rate of 1 mL/min in phosphate buffer saline using a Prominence high-pressure liquid chromatography (HPLC) system (Shimadzu). For molecular weight characterization, light scattering was measured with a miniDAWN multi-angle light scattering detector (Wyatt), connected to a RID-20A differential refractive index detector (Shimadzu) for quantitation of the total mass and to a SPD-20A UV detector (Shimadzu) for evaluation of the sole protein content. Chromatograms were collected and analyzed using the *ASTRA7* software (Wyatt), using an estimated *dn*/*dc* value of 0.185 ml/g). The calibration of the instrument was verified by injection of 10 μL of 2.5 mg/L monomeric BSA.

### Differential Scanning Fluorimetry (DSF)

DSF assays on recombinant GLT25D1/COLGALT1 samples (wild-type and mutants) at a concentration of 1 mg/mL in 25 mM HEPES/NaOH, 100 mM NaCl, pH 8.0 were performed using a Tycho NT.6 instrument (NanoTemper Technologies GmbH). Data were analyzed and plotted using *Graphpad Prism 7*^60^.

### Evaluation of Gal-T activity through luminescence

Reaction mixtures (5 μL total volume in 25 mM HEPES/NaOH, 100 mM NaCl, pH 8.0) were prepared by sequentially adding GLT25D1/COLGALT1 (final concentration 1 μM), 4 mg mL^-1^ of bovine skin gelatin, solubilized through heating denaturation at 37 °C for 10 min, 100 μM of UDP-α-sugar depending on the reaction, 50 μM MnCl_2_, and let incubate for 1 hour at 37 °C. The reactions were stopped by heating at 95 °C for 2 min, prior to transfer into Proxiplate white 384-well plates (Perkin-Elmer). At this point, 5 μL of the UDP-Glo luminescence detection reagent (Promega) were added and let incubate for 1 h at 25 °C. The plates were then transferred into a GloMax Discovery plate reader (Promega) configured according to manufacturer’s instructions for luminescence detection. All experiments were performed in triplicates. Control experiments were performed using identical conditions by selectively removing GLT25D1, donor or acceptor substrates. Data were analyzed and plotted using the *GraphPad Prism 7*^60^. Results of the data analysis are summarized in Supplementary Table 5.

### Evaluation of Gal-T activity through HR-LCMS

5 μM recombinant human LH3/PLOD3 (obtained in house as described in previous work^34^) was incubated with 5 μM GLT25D1/COLGALT1 (wild-type or mutants), 50 μM FeCl_2_, 100 μM 2-oxoglutarate (2-OG), 500 μM sodium ascorbate, 50 μM MnCl_2_, 100 μM UDP-α-sugar, and 1 mM peptide (Ac-GIKGIKGIKGIK-COOH) substrate. The reactions were allowed to proceed for 3 h at 37 °C. 5 μL of each reaction were diluted with 43 μL of Milli-Q water and acidified by addition of 2 μL of formic acid, to reach a final volume of 50 μL, and then analyzed on an Exion LC AD UHPLC (AB Sciex) coupled to a X500B ESI-HRMS/MS system (AB Sciex) by a full scan acquisition. The column oven was kept at 40 °C, while the autosampler was cooled at 10 °C. Peptides were separated by reverse phase HPLC on a Hypersil Gold C18 column (150 × 2.1 mm, 3 μm particle size, 175 Å pore size, Thermo Fisher Scientific) using a linear gradient (2-50% solvent B in 15 min), with solvent A consisting of 0.1% aqueous formic acid and solvent B of acetonitrile containing 0.1% formic acid. Flow rate was kept constant at 0.2 mL/min. Mass spectra were generated in positive polarity under constant instrumental conditions: ion spray voltage 4,500 V, declustering potential 100 V, curtain gas 30 psi, ion source gas 1 40 psi, ion source gas 2 45 psi, temperature 350 °C, collision energy 10 V. Spectra analyses were performed using the *SCIEX OS 2.1* software (AB Sciex). Results of the data analysis are summarized in Supplementary Table 5.

### Size Exclusion Chromatography coupled with Small-Angle X-ray Scattering (SEC-SAXS)

Solution scattering data were collected at ESRF BM29 using a sec^-1^ frame rate on Pilatus 11M detector located at a fixed distance of 2.871m from the sample, allowing a global *q* range of 0.01-4.001nm. SEC-SAXS experiments were carried out using Nexera High Pressure Liquid/Chromatography (Shimadzu) system connected online to SAXS sample capillary^61^. For these experiments, 50 μL of GTL25D1/COLGALT1 concentrated at 4 mg/mL were injected into a Superdex 200 PC 3.2/300 Increase column (GE Healthcare), pre-equilibrated with 25 mM HEPES/NaOH, 200 mM NaCl, pH 8.0. For SEC-SAXS data, frames corresponding to GTL25D1/COLGALT1 protein peak were identified, blank subtracted and averaged using *CHROMIXS*^62^.

### SEC-SAXS Modeling and data analysis

Radii of gyration (*Rg*), molar mass estimates and distance distribution functions *P(r)* were computed using *PRIMUS*^63^ within the *ATSAS* package^64^. Comparison of experimental SAXS data and 3D models from crystal structures was performed using *CRYSOL*^65^. A summary of SAXS data collection and analysis results is reported in Supplementary Table 6. *Ab initio* sphere models were generated from the SAXS data using *GASBOR*^66^, by providing the software with *P(r)* functions derived from experimental data and the number of residues defining GLT25D1/COLGALT1 monomers and without imposing any internal symmetry. 20 independent *ab initio GASBOR* runs were performed and subject to similarity analysis using *SUPCOMB* and *DAMSEL* from *ATSAS*^64^, allowing selection of a representative individual model based on normalized spatial discrepancy. Superpositions were carried out using the *SUPALM* program available in the *saspy ATSAS* plugin in *PyMol*^67^. For SAXS rigid-body modeling, the model of a GLT25D1/COLGALT1 monomer from the experimental crystal structure was subject to loop modeling using *CORAL*^68^. Evaluation of the agreement between molecular models and experimental SAXS profiles was carried out using *CRYSOL*^65^.

### Single particle negative staining electron microscopy

200 mesh, Cu grids (Ted Pella) grids were negatively charged using a PELCO EasiGlow (Ted Pella) with the following parameters: pressure 40 mBar, power 15 mA, time 30 s. 2 μL of 1 mg/mL GLT25D1/COLGALT1 were applied on each grid and left for 60 s. The excess sample was blotted using Whatman paper and then a 25 μl drop of 2% (w/v) uranyl acetate solution was applied, gently stirring for 1 min. The excess stain was blotted with filter paper and the grids were left dry overnight. 350 EM micrographs were acquired on a JEM 1200EXIII (JEOL) electron microscope equipped with a Mega View III CCD camera (Olympus). Data processing was carried out using *RELION* ^69^, including auto-picking and multiple rounds of 2D classification. Comparison of electron microscopy single particle analysis 2D classes and GLT25D1/COLGALT1 molecular models from crystal structures was carried out using *AlignProjections* within *the COSMIC2* webserver^70^.

## RESULTS

### GLT25D1/COLGALT1 is a Mn^2+^ dependent galactosyltransferase

We established methods to recombinantly produce and purify human full-length GLT25D1/COLGALT1 in *E. coli*and obtained a highly pure enzyme that could be used to evaluate its galactosyltransferase activity *in vitro*, through direct detection of Gal-T activity on synthetic collagen peptides using high-resolution mass spectrometry (HRMS), as well as by monitoring UDP generation from consumption of the UDP-α-Gal donor substrate in the presence of collagen hydroxylysines. Using these assays, we evaluated the enzyme’s strong preference for the Mn^2+^ metal ion cofactor, highlighting complete absence of enzymatic activity in presence of other divalent metal ions, with the exception of Mg^2+^, which nevertheless yielded much lower enzymatic activity compared to Mn^2+^ (Supplementary Fig. 1a). Given the ability of closely-related multifunctional LH/PLOD enzymes to process multiple donor substrates also in absence of acceptor substrates^33,34^, we probed GLT25D1/COLGALT1 and found that differently from LH/PLOD, GLT25D1/COLGALT1 is highly specific for UDP-α-Gal and displays very limited processing of UDP-α-Gal in absence of acceptor substrates (i.e., uncoupled activity, Supplementary Fig. 1b).

### The experimental crystal structure of human full-length GLT25D1/COLGALT1 reveals an elongated multi-domain architecture

The 2.8 Å crystal structure of human wild-type GLT25D1/COLGALT1 was obtained by experimental phasing, exploiting single wavelength anomalous dispersion (SAD) of a Hg^2+^ derivative (Supplementary Table 2). The quality of the electron density was excellent for the globular region of the two monomers of enzyme found in the asymmetric unit (Supplementary Fig. 2), whereas flexible regions located at the N-(residues 30-41) and C-termini (residues 572-618) could not be modeled.

Each GLT25D1/COLGALT1 polypeptide is characterized by two globular domains, which we named GT1 and GT2 (Fig. 1b), both characterized by similar glycosyltransferase architectures, but slightly different topological organization (Supplementary Fig. 3b). Although both structurally divergent from previously determined Rossmann fold architectures (the closest structure shows RMSD of 2.5 Å when superimposed), the GT1 domain appears more similar to known glycosyltransferases, yielding Z-scores higher than 15 with few bacterial and human glycosyltransferases (Supplementary Fig. 3b) according to DALI^71^. The closest homolog with an experimentally determined structure is human LH3/PLOD3^34^, whose central accessory (AC) domain shares approximately 25% sequence identity with GLT25D1/COLGALT1 GT1 domain and a remarkably similar structural arrangement with RMSD of 1.2 Å (Supplementary Fig. 3c). On the contrary, the GT2 domain shares only distant structural homology with known glycosyltransferases and the resulting superpositions appear much less clear compared to GT1 (Supplementary Fig. 3d). The two GT domains contact each other through electrostatics and hydrophobic contacts along helix α6 (GT1) and α7 (GT2). A 25-amino acid linker interconnects the two domains by shielding otherwise surface-exposed hydrophobic patches with two phenylalanine residues (Phe327 and Phe340, respectively, Fig. 1c).

### Both GT domains of GLT25D1/COLGALT1 can be populated with donor substrates and metal ion cofactors

Analysis of the experimental electron density of GLT25D1/COLGALT1 revealed intense difference peaks in the GT1 domain of both enzyme monomers proximate to the DxD motif identified by residues Asp166 and Asp168, in a loop interconnecting strand β4 and helix α4, that could unambiguously be modeled as Mn^2+^ and UDP-α-Gal (Fig. 2a). The carboxylate group of Asp166 is not directly involved in the coordination of the Mn^2+^ metal ion. The donor substrate is surrounded by an extended amino acid network, characterized by a H-bonded interaction involving the side chains of Tyr126 with the C^4^ carbonyl of the uridine moiety, which is trapped by a pi-Sigma interaction with Arg61 and a pi-Alkyl interaction with Val143, in a hydrophobic cavity also shaped by residues Leu59, Asp91, Arg139. The ribose moiety is kept into a well-defined conformation by Trp135, and by hydrogen bonds with the main chain carbonyl and amino groups of residues Leu59, Arg61, and Ala167. Additional interactions involve the side chains of Arg147 and Asp265 with hydroxyl groups of the galactose moiety (Fig. 2a,b). Notably, Trp135 matches Trp130 in the GLT25D1/COLGALT1 mouse protein sequence (Supplementary Fig. 4), whose mutation has been associated to skeletal and muscular defects due to severely reduced expression of the mutated gene^28^.

**Figure 2.**
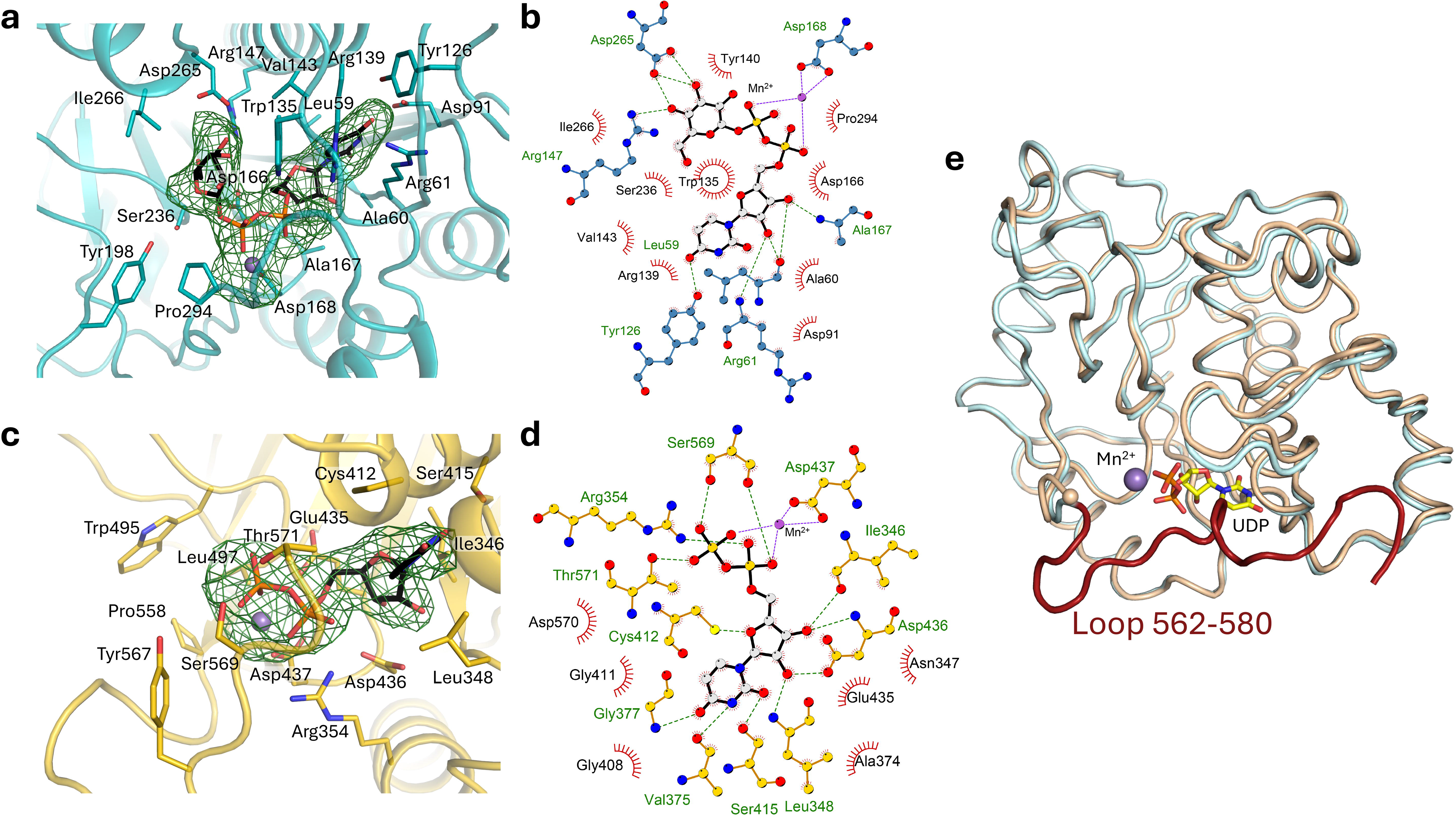
Structural insights into GLT25D1/COLGALT1 enzymatic activity and assembly. (a) The GT1 domain is constitutively populated by Mn^2+^ and UDP-α-Gal, as shown in the omit electron density map (contour level 3.2 σ). Residues involved in coordination of the metal ion and interaction with the donor substrate are shown as sticks; (b) LIGPLOT+^77^ diagram illustrating the interaction network for donor substrates and cofactors observed in the GT1 domain; (c) When co-crystallized with excess Mn^2+^ and UDP-α-Gal, also the electron density of the GT2 domain (omit map, contour level 3.2 σ) shows presence of donor substrates and cofactors. In this domain, the galactose moiety of the donor substrate is not visible, hence UDP has been modelled as sticks. Residues involved in coordination of the metal ion and interaction with the donor substrate are shown as sticks; (d) LIGPLOT+^77^ diagram illustrating the interaction network for donor substrates and cofactors observed in the GT2 domain; (e) Binding of Mn^2+^ and UDP-α-Gal induces conformational changes in the GT2 domain, by stabilizing the otherwise flexible C-terminal loop through direct interactions with the cofactor and the donor substrate. Shown is the superposition of GLT25D1 GT2 domains in substrate-free (light orange cartoon) *versus* substrate-bound (light blue cartoon) states, with highlight of the loop comprising residues 560-574 (brown), stabilized only when Mn^2+^ and UDP-α-Gal are present.

When co-crystallized with Mn^2+^ and UDP-α-Gal, the crystal structure of GLT25D1/COLGALT1 unexpectedly revealed electron density matching cofactors and donor substrates also in the GT2 domain of one of the two monomers in the asymmetric unit (Fig. 2c). In this domain the Mn^2+^ cofactor was found to an unusual Glu-Asp-Asp motif adopting a conformation topologically equivalent to that found in Mn^2+^-dependent glycosyltransferases, with the carboxyl groups of Glu435 and Asp437 directly chelating the metal ion, while the side chain of Asp436 was interacting through H-bonded interactions with the hydroxyl groups of the ribose moiety of the cofactor (Fig. 2c,d). The uridine ring of the donor substrate was constrained by Gly408 and Gly411 of α10-helix, in a hydrophobic groove defined by Cys412 of the same helix, plus Leu348 and Ala 374 on the opposite side (Fig. 2c,d). Notably, no electron density could be observed for the galactose moiety, suggesting multiple conformations for the sugar possibly due to the absence of acceptor substrates in the wide crevice defined by residues Trp495, Leu497, Tyr567, and Pro558 (Fig. 2c,d).

Comparison of GT2 domains with and without donor substrates highlighted a conformational rearrangement of the otherwise flexible C-terminus of the domain for residues 562-580, wrapping around the donor substrate binding cavity and positioning the backbone carbonyl of residue Ser569 and Thr571 in a productive conformation for hydrogen bonding with both phosphates of the UDP donor substrate (Fig. 2e). The presence of donor substrates and cofactors only in one of the two crystallographic monomers could be explained with the different crystal packing network surrounding the GLT25D1/COLGALT1 molecules, which prevents the rearrangement of the GT2 C-terminal loop in one of the two molecules due to crystal contacts (Supplementary Fig. 5).

### GLT25D1/COLGALT1 is a dimeric enzyme

The presence of two distinct GLT25D1/COLGALT1 molecules bound to different substrates and cofactors in the crystallographic asymmetric unit prompted us to investigate possible quaternary assemblies for this enzyme. The asymmetric unit of the GLT25D1/COLGALT1 crystal structure is indeed compatible with two enzyme molecules aligned in an antiparallel fashion through contacts mediated via their GT1 domain and wrapped around the unstructured N-terminal loop comprising residues 42-51 (Fig. 3a). This quaternary structure is characterized by a buried surface area of 1250 Å^2^, with 22 hydrogen bonds and 5 salt bridges out of 35 residues per monomer involved in the dimeric interface. Such assembly was confirmed by mass photometry (MP) analysis, showing that GLT25D1/COLGALT1 in solution is homogenously present with a molecular weight of 140 kDa, equivalent to the expected molecular weight for enzyme dimers (Fig. 3b). Consistent results of 131 ± 10 kDa were obtained using size exclusion chromatography coupled to multi-angle light scattering (SEC-MALS) and small angle X-ray scattering (SEC-SAXS) (Supplementary Table 6, Supplementary Fig. 6), the latter producing *ab initio* sphere models of elongated molecular ensembles matching the size and overall envelope shape of crystallographic GLT25D1/COLGALT1 dimers with 95% correlation. Orthogonal comparison of the unmodified experimental crystal structure with solution SAXS data yielded initial *χ*^2^ of 7.62, which further improved to 1.89 after modeling of flexible N- and C-termini (Fig. 3c). Furthermore, low-resolution 2D classes from a sample of GLT25D1/COLGALT1 analyzed using single particle negative staining electron microscopy revealed elongated objects highly comparable to the SEC-SAXS solution data and the crystallographic dimers (Fig. 3d).

**Figure 3.**
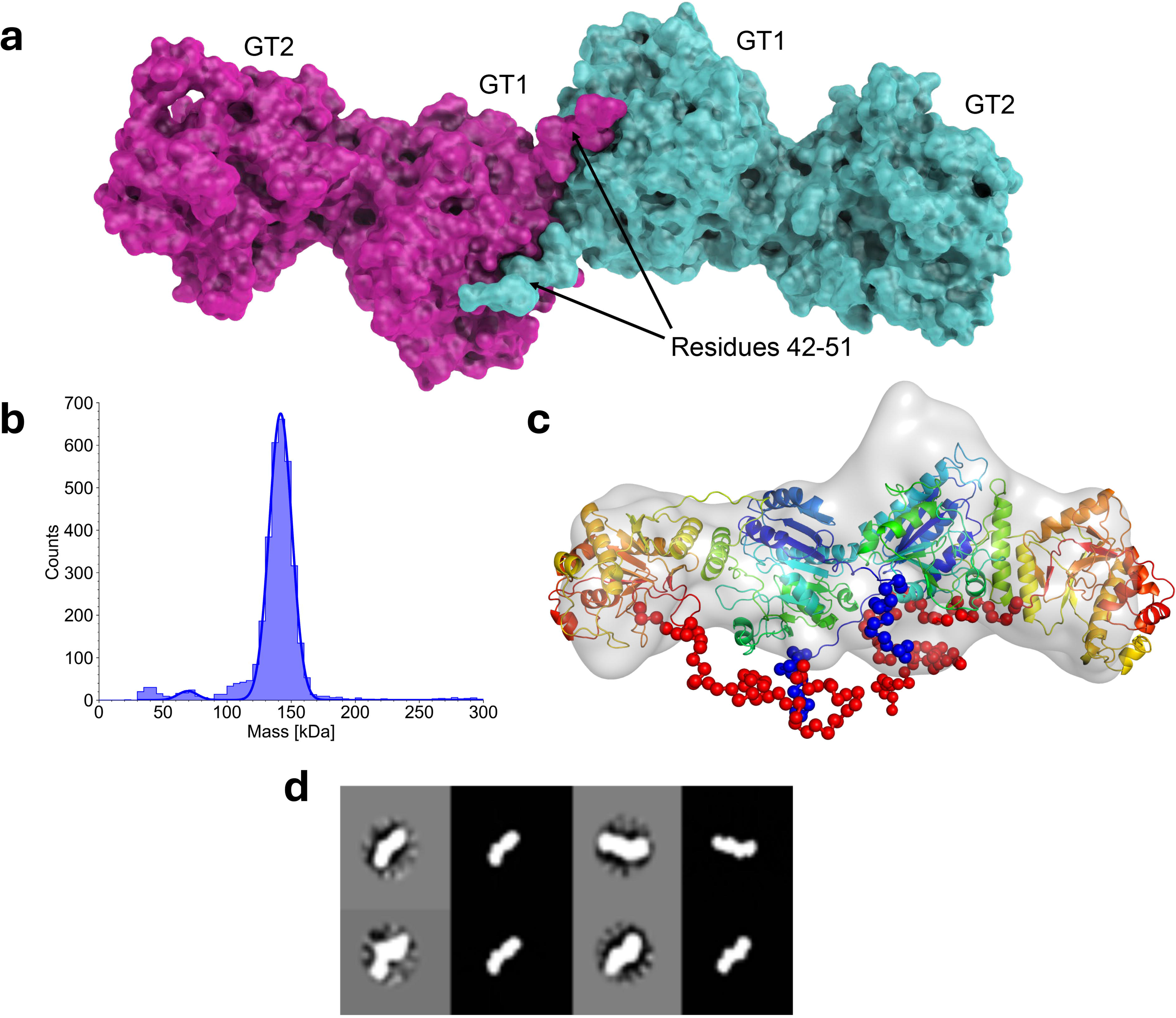
GLT25D1/COLGALT1 forms stable elongated dimers. (a) Structure of dimeric full-length GLT25D1/COLGALT1 enzyme, showing two molecules interconnected through their N-terminal domains. (b) Mass photometry analysis of recombinant wild-type GLT25D1/COLGALT1 samples; (c) Modeling of the N- and C-terminal flexible loops (shown as spheres) of GLT25D1/COLGALT1 yields a structural model of the dimeric enzyme (colored blue to red from the N- to the C-terminus of each monomer) that fits the experimental solution SAXS data with *χ*^2^ of 1.89, superimposed to the envelope surface computed from *Ab-initio* sphere model based on solution SEC-SAXS data match the observed dimeric arrangement found in the experimental crystal structure of GLT25D1/COLGALT1; (d) Comparison of NS EM SPA 2D classes (gray background) with selected orientations of computed projections of the GLT25D1/COLGALT1 crystal structure (black background) shows matching molecular projections.

### Dimerization of GLT25D1/COLGALT1 is not essential for enzyme function

To evaluate the importance of dimerization on GLT25D1/COLGALT1 function, we designed point mutations for amino acid residues located at the crystallographic dimer interface: Trp158, whose side chain is embedded into a cavity shaped by Arg53, Ala85, His113, Met157 of the other enzyme monomer; and Asp160, forming a hydrogen bonding network with Arg53 from both monomers (Fig. 4a). Mutation Trp158Arg was enough to disrupt the dimer interface of GLT25D1, resulting in monomeric species in solution as assessed by mass photometry and SEC-MALS (Fig. 4b,c, Supplementary Fig. 7). When prompted for enzymatic activity using either indirect luminescence-based detection of free UDP in the presence of gelatin, as well as direct evaluation of galactosyl-hydroyxlysine modification of synthetic peptides using high-resolution mass spectrometry (HRMS), this mutant showed activities comparable to wild-type GLT25D1/COLGALT1 (Fig. 4d, Supplementary Table 5), supporting the hypothesis that enzyme dimerization does not constitute an essential requirement for enzyme function *in vitro*.

**Figure 4.**
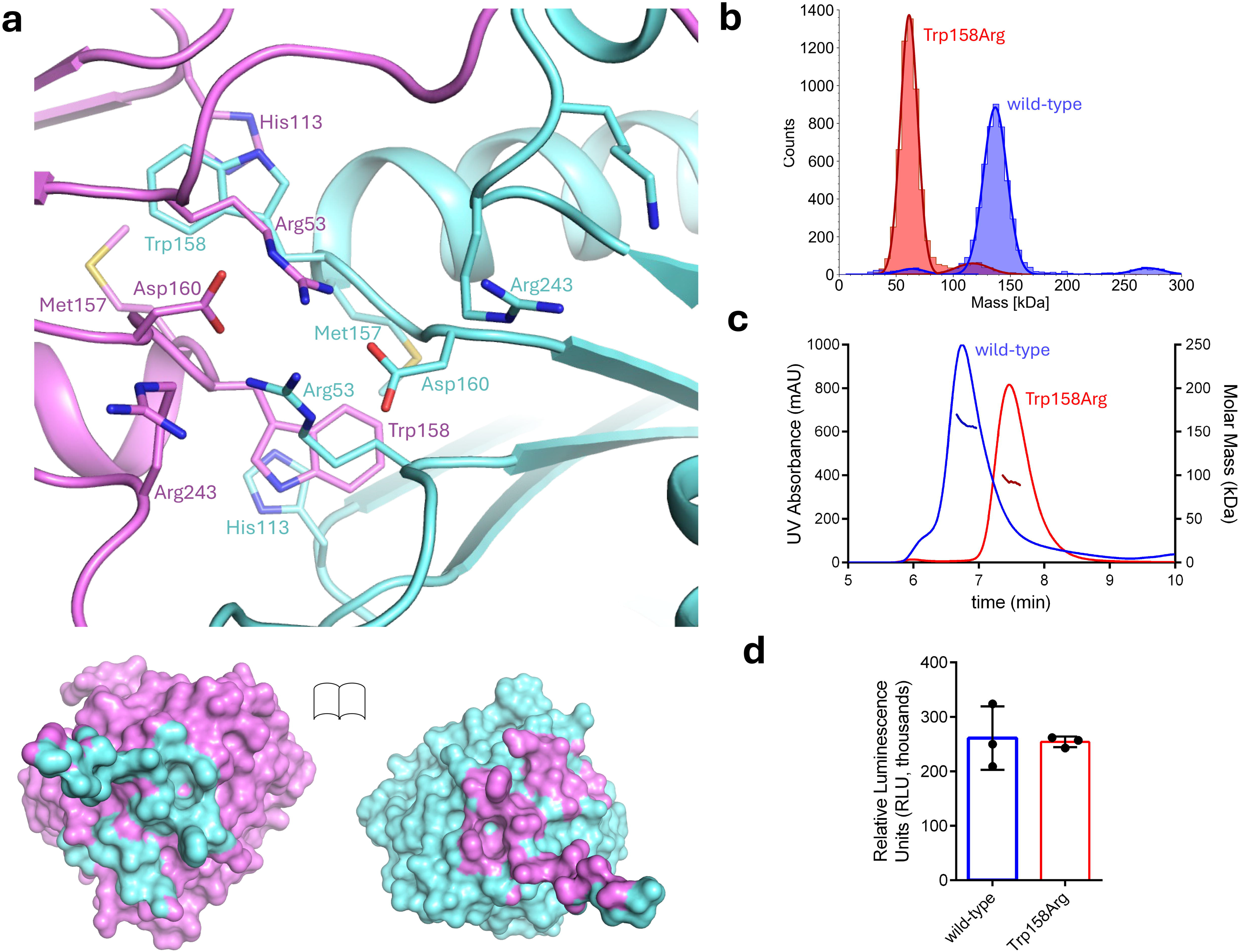
Probing GLT25D1/COLGALT1 dimeric architecture and its functional significance. (a) Details of the dimeric interface, showing amino acid residues involved in the direct contacts between the two monomers. The bottom panel shows an open-book rendering of the two GLT25D1/COLGALT1 monomers, with the colored footprint of the dimerization partner on the molecular surface; (b) Results of the mass photometry comparison of wild-type GLT25D1/COLGALT1 and the Trp158Arg mutant, designed to disrupt the dimer interface; (c) Results of the SEC-MALS comparison of wild-type GLT25D1/COLGALT1 and the Trp158Arg mutant, showing alterations of the elution volume as well as molar mass; (d) Comparison of the Gal-T enzymatic activity of GLT25D1/COLGALT1 wild-type and Trp158Arg mutant using luminescence. The histograms show the average result of triplicate independent measurements (whose results are individually shown as overlayed dots). Error bars represent standard deviations from average of triplicate independent experiments.

### Cofactors and donor substrates in GLT25D1/COLGALT1 GT1 domain have structural roles

Intrigued by the observation of a dimeric ensemble potentially characterized by four aligned catalytic sites, we wondered whether metal ions and donor substrates could have structural, rather than functional roles in the GT1 domain. We therefore purified recombinant GLT25D1/COLGALT1 without any Mn^2+^ supplementation and attempted to remove the tightly bound Mn^2+^ by incubating the enzyme with different concentrations of ethylenediaminetetraacetic Acid (EDTA). We consistently observed a destabilization, with a differential scanning fluorimetry (DSF) thermal shift of 4.7 °C compared to untreated samples (Fig. 5a). Nevertheless, EDTA-treated samples still showed DSF profiles compatible with a folded protein sample, thus we wondered whether removal of metal ions through EDTA treatment could impact on the enzyme’s oligomeric state. EDTA-treated GLT25D1/COLGALT1 samples were therefore subject to size exclusion chromatography analysis. Stripping of the metal ion altered the elution profile of the enzyme, yielding heterogenous oligomeric species (Fig. 5b).

**Figure 5.**
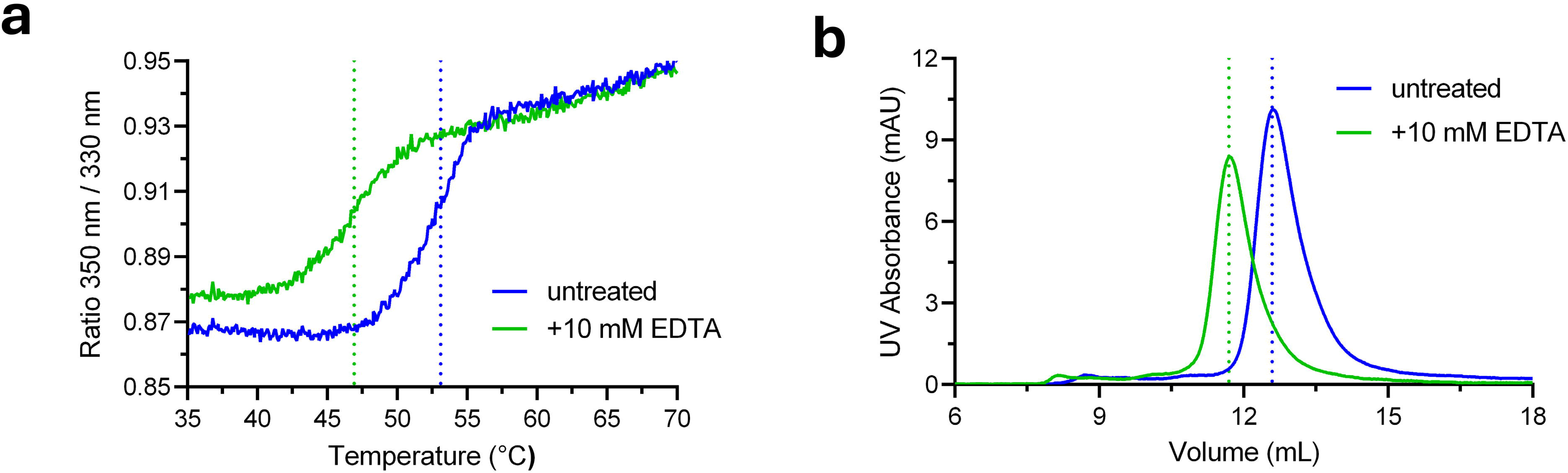
Probing the structural role of Mn. ^2+^ **in GLT25D1/COLGALT1 GT1 domain.** (a) DSF analysis of GLT25D1/COLGALT1 subject to EDTA treatment, showing the destabilization induced by Mn^2+^ chelation; (b) EDTA-treated samples show altered elution profiles when tested using size-exclusion chromatography.

Structure-guided point mutants designed to interfere with Mn^2+^ or UDP-α-Gal binding in the GT1 domain (Fig. 2a,b) such as Asp166Ala (abolishing interaction with Mn^2+^), Asp265Ala (removing H-bonding contacts with the galactose moiety), Ile266Gln (preventing rigid positioning of the galactose moiety in the GT1 cavity), did not produce any recombinant enzyme samples suitable for biochemical studies, and were consistent with previously published attempts describing a Asp166Ala-Asp168Ala double mutant, designed to abolish the GT1 DxD motif^37^. Control mutants such as Ile267Ala (in close proximity to Ile266, but not affecting UDP-α-Gal binding or positioning) instead resulted in well folded and active enzyme samples (Supplementary Fig. 8), matching the previously published Pro292Asn mutant, originally designed to generate a chimeric GLT25D1-CERCAM inactive galactosyltransferase, but still retaining enzymatic activity^37^. Collectively, these data support structural roles for metal ion cofactors and donor substrates in the GT1 domain without direct functional roles during catalysis.

### The GLT25D1/COLGALT1 GT2 domain is responsible for the catalytic activity

Our experimental structure investigations revealed that the GT2 domain of GLT25D1/COLGALT1 binds to the Mn^2+^ cofactor and donor substrates transiently compared to GT1, inducing conformational changes of the extended C-terminus of the enzyme that could have an impact on catalysis (Fig. 2). The identification of an unprecedented EDD motif coordinating Mn^2+^ within this domain, together with the presence of a UDP-α-Gal donor substrate with a flexible sugar moiety (Fig. 2c,d), may provide ultimate insights for the identification of the enzyme’s catalytic site. Conservation maps for the amino acid residues shaping the two putative ligand binding sites in homologous vertebrate enzymes highlighted that the amino acids involved in Mn^2+^ and UDP-α-Gal binding are highly conserved in both GT1 and GT2 domains (Fig. 6a). Prompted by this observation, and by our insights suggesting structural roles for the enzyme’s N-terminal GT1 domain, we generated a series of point mutations affecting Mn^2+^ and donor substrate binding in the GT2 domain, as well as residues potentially involved in acceptor substrate binding during transfer of the galactose moiety. All mutants generated could be purified and showed biophysical features compatible with well-folded proteins (Supplementary Fig. 9). Mutant Asp437Ala, designed to abolish Mn^2+^ coordination, was completely inactive in both luminescence and HRMS assays (Fig. 6b, Supplementary Table 5). Similar results could be obtained with mutant Cys412Ser, affecting productive positioning of the uridine-ribose moieties in a hydrophobic cavity. Inspired by the potential roles found for aromatic and charged residues in the glycosyltransferase enzyme cavity of LH/PLOD enzymes^33^, we generated mutants Trp495Ala and Asp522Ala, which both resulted in complete loss of GLT25D1/COLGALT1 Gal-T activity (Fig. 6b, Supplementary Table 5). The structural data obtained also provided a rationale for previously investigated mutations^37^: Asp336, located at the C-terminus of the linker segment connecting GT1 and GT2 domains, whose hydrogen bond with GT2 Lys445 does not seem to be dramatically impacted when being mutated into a serine residue; and residues Asp461 and Asp433, the latter being involved in a hydrogen bonding network with the side chains of Lys508 and His547, likely critical for GT2 domain folding stability. Taken together, our structure-guided site-directed mutagenesis results enabled the unambiguous identification of GLT25D1/COLGALT1 catalytic site and its unusual features.

**Figure 6.**
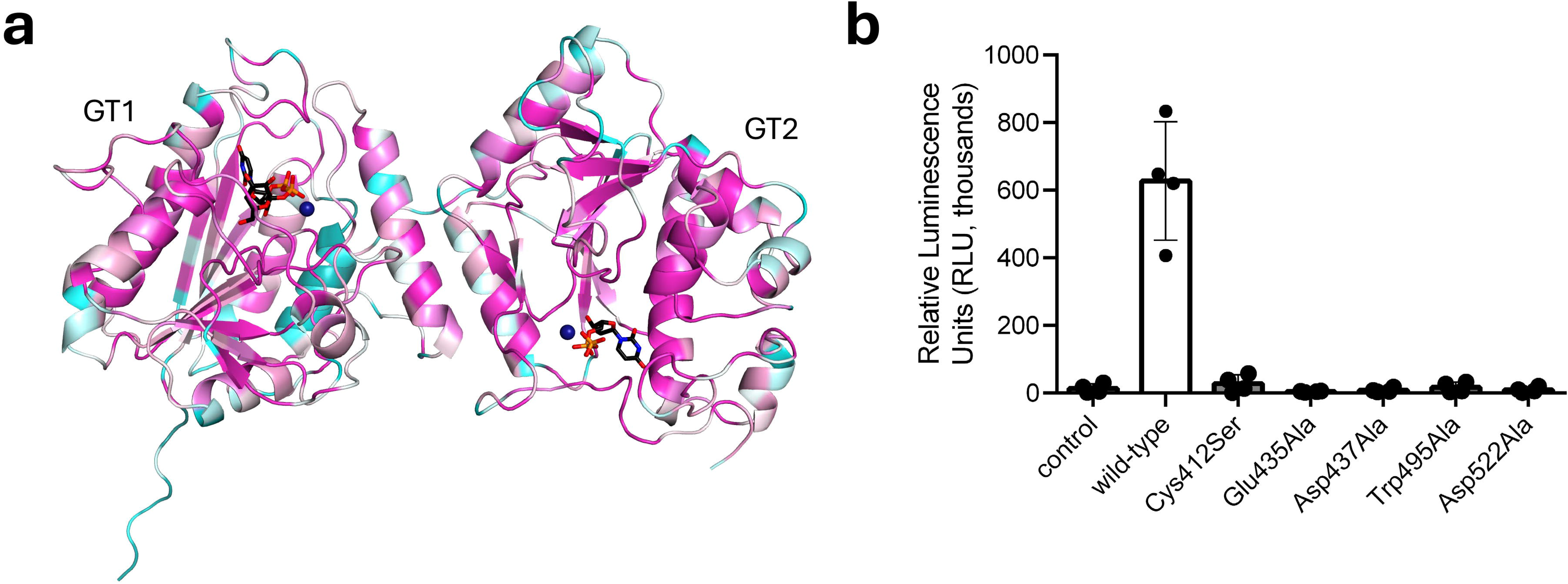
Individual GT domains are not sufficient for GLT25D1/COLGALT1 enzymatic activity. (a) Amino acid conservation diagram for different GLT25D/COLGALT homologs. The colors represent the different degree of amino acid conservation from dark blue (not conserved at all) to dark purple (fully conserved). The complete list used for the representation and the associated alignment is shown in Supplementary Fig. 4; (b) Luminescence-based comparison of the Gal-T enzymatic activity of wild-type GLT25D1/COLGALT1 and mutants designed to interfere with cofactor and substrate binding in the GT2 domain. The histograms show the average result of triplicate independent measurements (whose results are individually shown as overlayed dots). Error bars represent standard deviations from average of triplicate independent experiments.

## DISCUSSION

Collagen hydroxylysine galactosylation is a key step in the complex post-translational modification pathway that leads to Glc-Gal-Hyl, the most abundant O-linked glycosylation found in the animal kingdom^12,13,16^. Following decades of intensive biochemical investigations addressing the enzymes involved in this process and their associated functions, the recent years have witnessed increasing efforts to produce a comprehensive molecular description of these reactions and the actors involved, with particular attention to multifunctional LH/PLOD lysyl hydroxylases-glucosyltransferases^17,34^. The lack of experimental structure data about GLT25D/COLGALT galactosyltransferases have limited our understanding of the molecular mechanisms of these glycosyltransferases, despite previous significant efforts^16,27,37^. In this work, we have determined the experimental structure of full-length human GLT25D1/COLGALT1, and used a combination of biochemistry, biophysics and site-directed mutagenesis to characterize its functionality *in vitro*.

Our results highlighted an elongated, multi-domain dimeric quaternary structure characterized by two Rossmann fold-type domains per monomer (named GT1 and GT2, respectively), interconnected through a linker region characterized by a rigid conformation and with dimer contacts wrapping around the N-terminus of each monomer (Fig. 1b,c). Surprisingly, both GT1 and GT2 domains were indispensable for catalysis and were found populated by the Mn^2+^ cofactor as well as by the UDP-α-Gal donor substrate (Fig. 2), with the ligand binding regions of both domains showing the highest degree of sequence conservation (Fig. 5a, Supplementary Fig. 4). The N-terminal domain GT1 is closely related to the inactive central accessory (AC) domain of LH/PLOD enzymes^34^ and shares features more similar those typical of inverting GT-A glycosyltransferases. Conversely, the GT2 domain shares much less similarity with known glycosyltransferases, but at the same time displays higher amino acid sequence conservation for the residues surrounding the site occupied by the Mn^2+^ cofactor and UDP-α-Gal (Fig. 6a, Supplementary Fig. 4). Previous work addressing the putative functional roles of GLT25D1/COLGALT1 DxD motifs showed that point mutations affecting residues involved in Mn^2+^ coordination Asp166 and Asp168 in the GT1 domain completely abolished the activity^37^. This is consistent with our identification of Mn^2+^ and UDP-α-Gal bound in the GT1 domain of the enzyme. However, the same authors reported that replacing the entire N-terminus of the enzyme with that of inactive CERCAM did not impact on the ability of the enzyme to glycosylate collagen substrates^37^. Such apparently contradictory result can be explained with the unprecedented structural role of Mn^2+^ and UDP-α-Gal in this domain, both essential to preserve the folding stability of GT1 and its dimeric quaternary structure, but not required for a galactosyltransferase enzymatic activity which, as we showed with site-directed mutagenesis experiments, does not depend on enzyme dimerization (Fig. 4d).

Analogously, previous attempts to map the catalytic residues critical for GLT25D1/COLGALT1 enzymatic activity were not conclusive^37^, likely due to the unexpected unique Glu-Asp-Asp motif that we found responsible for binding Mn^2+^ in the GT2 domain, and the multiple DxD motifs within the enzyme polypeptide sequence (Supplementary Fig. 4). To ensure correct mapping of the GT2 catalytic network, we therefore designed several point mutations surrounding the Mn^2+^ cofactor and the donor substrate and found an extensive network of charged and hydrophobic side chains responsible for UDP-α-Gal substrate activation and catalysis (Fig. 6b).

The overall arrangement of substrate-free and substrate-bound conformations for the GT2 catalytic cavity is almost identical (Figure 2), except for the conformational rearrangement of the otherwise flexible C-terminal segment of the domain. This region is characterized by negatively charged amino acids, whose direct involvement in binding and processing of the donor substrate UDP-α-Gal has been demonstrated by our site-directed mutagenesis experiments. In this respect, the new experimental insights also support previous *in silico* hypotheses^72^.

The observed dimeric quaternary structure, confirmed by multiple orthogonal experiments in solution and by site-directed mutagenesis, does not appear to be a requirement for GLT25D1/COLGALT1 enzymatic activity *in vitro*. We speculate that the observed dimeric architecture could be a requirement for processing the complex collagen substrates *in vivo*, likely through molecular interactions with other collagen biosynthesis enzymes and/or chaperones. Of note, GLT25D/COLGALT enzymes have been proposed to directly interact with partners LH/PLOD in the endoplasmic reticulum, possibly forming biosynthetic complexes capable of fully converting collagen Lys residues into Glc-Gal-Hyl^26,45^. Interestingly, the unusually elongated shape of the GLT25D1/COLGALT1 dimer resembles that of the tail-to-tail dimer observed for LH3/PLOD3^34^, possibly facilitating extensive interactions among the two enzymes responsible for the subsequent processing of collagen lysines into Glc-Gal-Hyl and posing questions regarding the possible evolution of this two-enzyme system from a common ancestor. This is further supported by the structural similarity observed between GLT25D1/COLGALT1 GT1 domain and the LH/PLOD AC domain (Supplementary Fig. 3). Such hypothesis prompts for extensive future work, also taking into account the unusual trafficking of multifunctional LH/PLOD enzymes through the secretory pathway due to absence of specific ER retention sequences^73–76^.

## ACKNOWLEDGEMENTS

We thank Ms. Lisa Negro and Ms. Martina Soffientini for assistance with production of recombinant GLT25D1/COLGALT1, Dr. Marco Fumagalli for support with HRMS assays, and Dr. Gianluca Santoni for support with SAD X-ray data collection setup. We thank the European Synchrotron Radiation Facility (ESRF) and the Swiss Light Source (SLS) for the provision of synchrotron radiation facilities. We thank Centro Grandi Strumenti (University of Pavia) for the provision of HRMS and NS EM instrumentation.

## AUTHOR CONTRIBUTIONS

FF conceived the project and designed research. MDM, LS, SRR produced recombinant GLT25D1/COLGALT1 and performed biochemical characterizations, with support from DM. LS crystallized GLT25D1/COLGALT1. FF and LS solved the structure and carried out structural refinements, with support from MDM. DM and MDM performed HRMS assays. SL carried out mass photometry data analysis. AP generated all mutants. MDM, LS, and FF analyzed the data, prepared the figures and wrote the paper, with contributions from all authors.

## DATA AVAILABILITY

Coordinates and structure factors have been deposited in the Protein Data Bank under accession codes 9EVJ, 9EVK, and 9EVL. In solution SEC-SAXS data have been deposited in the SASBDB under accession code SASXXXX. All data needed to evaluate the conclusions in the paper are present in the paper or in the Supplementary Materials.

