## Supplementary Information for "Molecular Structure and Enzymatic Mechanism of the Human Collagen Hydroxylysine Galactosyltransferase GLT25D1/COLGALT1"

**Supplementary Table 1. List of oligonucleotides used to generate GLT25D1/COLGALT1 mutants**

| Site | Mutation | Direction | Sequence (5'→3') |
|---|---|---|---|
| GT1, dimer interface | Trp158Arg | Forward | ggctgattacatcctgtttgtagat |
| | | Reverse | CTaccgagacctcagtcgtatggaacaat |
| GT1, cofactor binding site | Asp166Ala | Forward | CAgcggacaacctgatcctcaaccc |
| | | Reverse | ctacaaacaggatgtaatcagcccacatgt |
| | Asp265Ala | Forward | catcatcgtctttgccttctcctgcaag |
| | | Reverse | Gcgtcaaaggaccaggtgtagtcagggt |
| | Ile266Gln | Forward | CAAatcgtctttgccttctcctgcaag |
| | | Reverse | gtcgtcaaaggaccaggtgtagtcaggg |
| | Ile267Gln | Forward | CAAgtctttgccttctcctgcaagcagg |
| | | Reverse | gatgtcgtcaaaggaccaggtgtagt |
| GT2, cofactor binding site | Cys412Ser | Forward | Gcttcctgagccactacaacatctggaag |
| | | Reverse | cttccagatgttgtagtggctcaggaagc |
| | Glu435Ala | Forward | ggatgacctgcgttttgagatcttcttc |
| | | Reverse | Gcaaacacaagcgatttctgcagcccc |
| | Asp437Ala | Forward | Ccctgcgttttgagatcttcttcaagagac |
| | | Reverse | catcctcaaacacaagcgatttctgcag |
| | Trp495Ala | Forward | gaccctggcctacgtgatctccctgcaag |
| | | Reverse | GCgtaggaatagtcagcttcgactaggtt |
| | Asp522Ala | Forward | cgagttcctgcccgtcatgttcgacaaac |
| | | Reverse | Gctactggaagcattttggagagcggc |

**Supplementary Table 2. X-ray diffraction data collection parameters and statistics**

| | GLT25D1/COLGALT1 (Native) | GLT25D1/COLGALT1 (co-crystallized with $Mn^{2+}$, UDP-α-Gal) | GLT25D1/COLGALT1 ($Hg^{2+}$ Soak) |
|---|---|---|---|
| **Data Collection**[a] | | | |
| X-ray source | ESRF ID23-1 | SLS X06SA | ESRF ID23-1 |
| Processing programs | *XDS*, *AIMLESS* | *XDS*, *AIMLESS* | *XDS*, *AIMLESS*, *SHELX*, *HKL2MAP* |
| Space group | I23 | I23 | I23 |
| Cell parameters | *a* = 221.9 Å *a* = 90.0°<br>*b* = 221.9 Å *b* = 90.0°<br>*c* = 221.9 Å *g* = 90.0° | *a* = 220.8 Å *a* = 90.0°<br>*b* = 220.8 Å *b* = 90.0°<br>*c* = 220.8 Å *g* = 90.0° | *a* = 220.2 Å *a* = 90.0°<br>*b* = 220.2 Å *b* = 90.0°<br>*c* = 220.2 Å *g* = 90.0° |
| Wavelength (Å) | 0.8856 | 0.9999 | 0.7749 |
| Resolution range (Å) | 49.62-3.00 (3.13-3.00) | 49.38-2.70 (2.79-2.70) | 49.24-2.80 (2.91-2.80) |
| Total reflections | 123628 (15938) | 425956 (23149) | 1013021 (105836) |
| Unique reflections | 35735 (4385) | 49018 (4444) | 43652 (4551) |
| CC1/2[b] | 0.992 (0.402) | 0.998 (0.395) | 0.998 (0.599) |
| Redundancy | 3.5 (3.6) | 8.7 (5.2) | 23.2 (23.3) |
| Mean I/σ(I) | 6.2 (1.0) | 13.5 (1.1) | 13.2 (0.9) |
| Completeness (%) | 98.4 (99.9) | 99.7 (97.9) | 100.0 (100.0) |
| $R_{sym}$[b] | 0.130 (1.047) | 0.110 (1.251) | 0.169 (3.609) |
| $R_{pim}$[c] | 0.101 (0.811) | 0.058 (0.920) | 0.051 (1.093) |
| Anomalous completeness (%) | | | 100.0 (100.0 |
| Anomalous multiplicity | | | 11.7 (11.7) |
| Anomalous CC1/2 | | | 0.488 |
| SAD resolution limit (Å)[d] | | | 4.4 |

[a] Values in parentheses are for reflections in the highest resolution shell.

[b] $R_{sym} = [\ \Sigma_{hkl}\Sigma_j\ |\ I_{hkl,j} - \langle I_{hkl}\rangle\ |\ ]\ /\ [\ \Sigma_{hkl}\Sigma_j\ I_{hkl,j}\ ]$ , where I is the observed intensity for a reflection and <I> is the average intensity obtained from multiple observations of symmetry-related reflections.

[c] $R_{pim} = [\ \Sigma_{hkl}\,(1/(n-1))^{1/2}\ \Sigma_j\ |\ I_{hkl,j} - \langle I_{hkl}\rangle\ |\ ]\ /\ [\ \Sigma_{hkl}\Sigma_j\ I_{hkl,j}]$ where I is the observed intensity for a reflection and <I> is the average intensity obtained from multiple observations of symmetry-related reflections.

[d] the resolution limit for SAD phasing has been selected by evaluating the resolution where *d''/s* falls below 1.2 using the *SHELXC* module of *HKL2MAP*.

**Supplementary Table 3. X-ray crystallographic refinement statistics**

| | GLT25D1/COLGALT1 (Native) | GLT25D1/COLGALT1 (co-crystallized with $Mn^{2+}$, UDP-a-Gal) | GLT25D1/COLGALT1 ($Hg^{2+}$ Soak) |
|---|---|---|---|
| **Refinement** | | | |
| $R_{work}/R_{free}{}^{c}$ | 0.1928/0.2392 | 0.1856/0.2220 | 0.2070/0.2428 |
| Number of atoms: | 8633 | 8920 | 8620 |
| Protein | 8519 | 8681 | 8503 |
| Ligands | 105 | 106 | 114 |
| Solvent | 9 | 133 | 3 |
| Average B-factor $(Å)^2$ | 91.20 | 74.89 | 109.26 |
| Protein | 91.37 | 75.22 | 109.14 |
| Ligands | 79.40 | 65.03 | 119.08 |
| Solvent | 64.04 | 54.24 | 78.75 |
| **Structure quality** | | | |
| RMS bond lengths (Å) | 0.003 | 0.004 | 0.002 |
| RMS bond angles (°) | 0.65 | 0.95 | 0.55 |
| **Ramachandran stats** | | | |
| Favored (%) | 95.84 | 97.05 | 95.36 |
| allowed (%) | 4.16 | 2.86 | 4.64 |
| outliers (%) | 0.00 | 0.10 | 0.00 |
| PDB ID | 9EVK | 9EVJ | 9EVL |

**Supplementary Table 4. Details of mass photometry analysis**

| GLT25D1/COLGALT1 Sample | Shown in | Mass (kDa) | Intensity Counts | Peak mass fraction |
|---|---|---|---|---|
| wild type | Figure 3B | 142.0 ± 8.8 | 2991 | 85% |
| wild type | Figure 4B | 137.0 ± 10.4 | 4597 | 83% |
| Trp158Arg | Figure 4B | 61.0 ± 7.6 | 5213 | 88% |

**Supplementary Table 5. Summary of the outcome of the luminescence-based indirect activity assays, and the HRMS direct assays to evaluate GLT25D1/COLGALT1 enzymatic activity.** Values are reported as percentage of enzymatic activity observed for each variant compared to that observed using the wild-type enzyme. Values are expressed as average of minimum triplicate independent measurements, with associated standard deviations.

| GLT25D1/COLGALT1 variant | Gal-T activity (%) | |
|---|---|---|
| | Luminescence-based indirect assay | HRMS direct assay |
| wild-type | 100.0 ± 13.4 | 100.0 ± 22.1 |
| Trp158Arg | 66.0 ± 6.3 | 97.2 ± 4.3 |
| Ile267Gln | 73.0 ± 6.2 | 93.3 ± 8.9 |
| Cys412Ser | 1.3 ± 1.5 | n.d. |
| Glu435Ala | n.d. | n.d. |
| Asp437Ala | n.d. | n.d. |
| Trp495Ala | n.d. | n.d. |
| Asp522Ala | n.d. | n.d. |

**Supplementary Table 6. SEC-SAXS data collection parameters and processing statistics**

| | |
|---|---|
| **Data Collection** | |
| Beamline | ESRF BM29 |
| Beam energy (keV) | 12.5 |
| Sample-detector distance (m) | 2.867 |
| Exposure time (s) | 1 |
| Sample cell thickness (mm) | 1 |
| Temperature (°) | 20 |
| Final q range ($nm^{-1}$) | 0.01 – 4.00 |
| **Data Analysis** | |
| Points used for Guinier analysis | 10 - 33 |
| Guinier $qR_g$ limits | 0.77 - 1.28 |
| Guinier $R_g$ (nm) | 4.33 ± 0.02 |
| I(0) ($mm^{-1}$) | 22.91 ± 0.08 |
| $D_{max}$ (nm) | 13.97 |
| MW estimation (Bayesian Inference) (kDa) | 131 ± 10 |

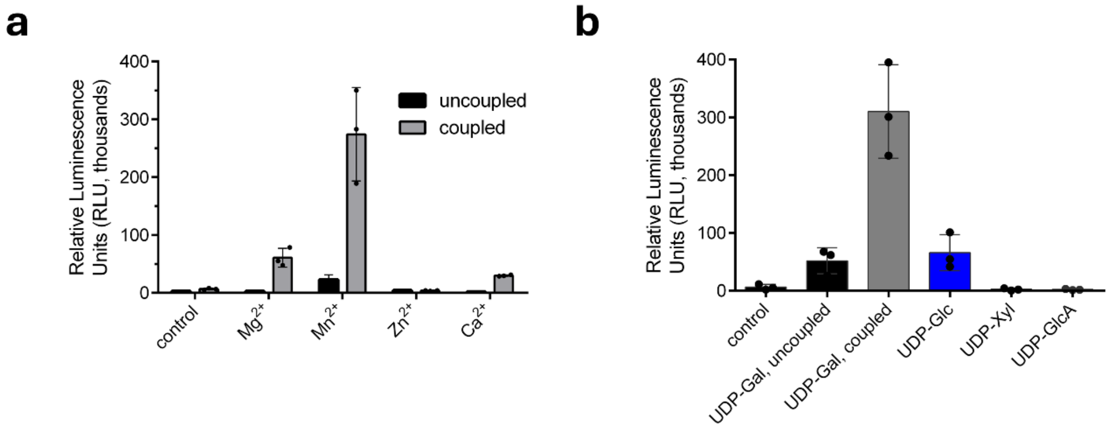


**Supplementary Figure 1. GLT25D1/COLGALT1 is a $Mn^{2+}$-dependent galactosyltransferase.** Luminescence-based assays to evaluate the galactosyltransferase enzymatic activity in presence of different metal ions (a) and different donor substrates and their analogs (b). Control experiments were performed without adding GLT25D1/COLGALT1. Uncoupled measurements were carried out in absence of acceptor (i.e., gelatin) substrate.

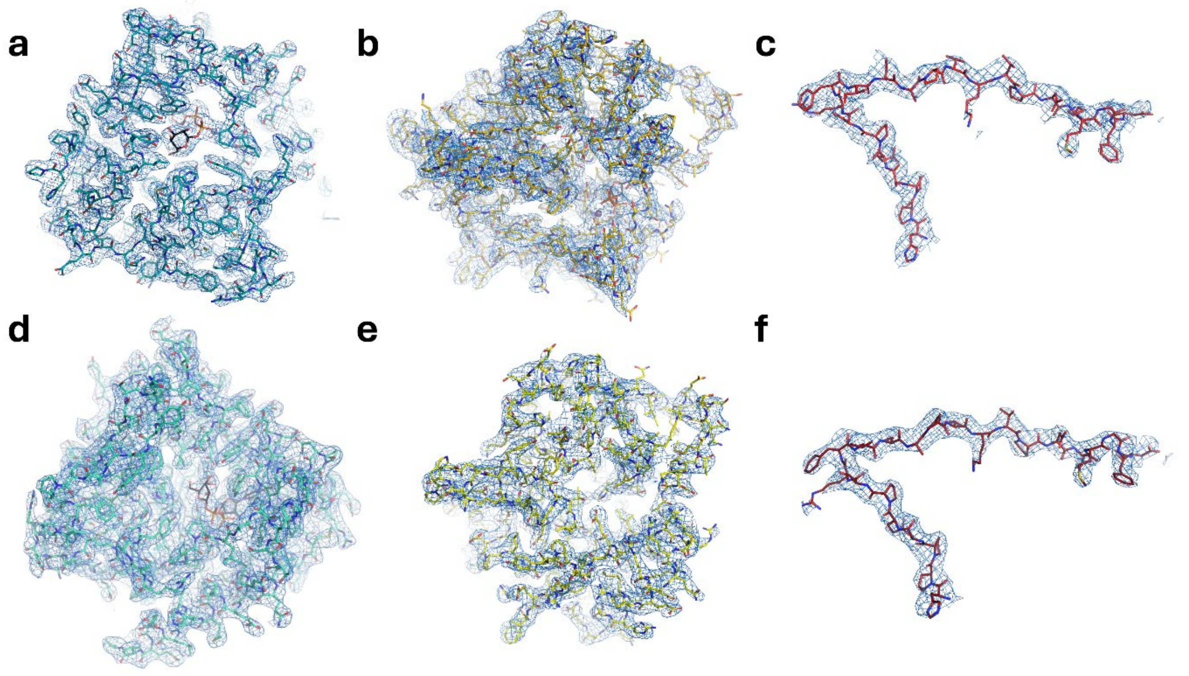


**Supplementary Figure 2. Quality of the experimental electron density for the crystal structure of full-length human GLT25D1/COLGALT1.** Shown are snapshots of the refined experimental electron density ($2F_o$-$F_c$, contour level 1 $\sigma$) used to generate the GLT25D1/COLGALT1 structural model using the $Mn^{2+}$, UDP-$\alpha$-Gal co-crystallized model, covering the GT1 domains (a,d), the the GT2 domains (b,e), and the linker regions (c,f) for each of the two copies of the enzyme found in the asymmetric unit.

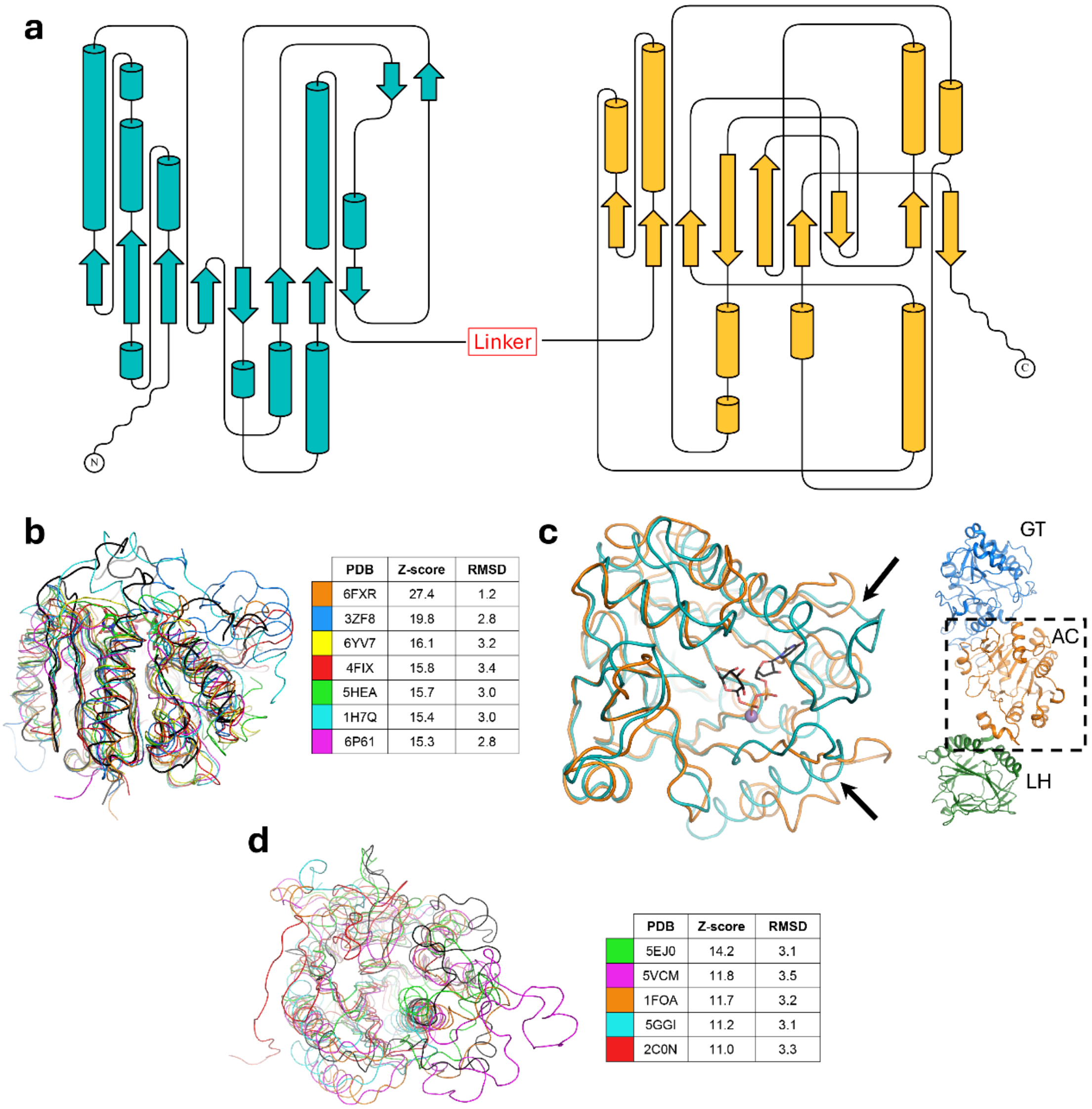


| | PDB | Z-score | RMSD |
|---|---|---|---|
| (orange) | 6FXR | 27.4 | 1.2 |
| (blue) | 3ZF8 | 19.8 | 2.8 |
| (yellow) | 6YV7 | 16.1 | 3.2 |
| (red) | 4FIX | 15.8 | 3.4 |
| (green) | 5HEA | 15.7 | 3.0 |
| (cyan) | 1H7Q | 15.4 | 3.0 |
| (magenta) | 6P61 | 15.3 | 2.8 |

| | PDB | Z-score | RMSD |
|---|---|---|---|
| (green) | 5EJ0 | 14.2 | 3.1 |
| (magenta) | 5VCM | 11.8 | 3.5 |
| (orange) | 1FOA | 11.7 | 3.2 |
| (cyan) | 5GGI | 11.2 | 3.1 |
| (red) | 2C0N | 11.0 | 3.3 |

**Supplementary Figure 3. Topological details of GLT25D1/COLGALT1 GT1 and GT2 domains.** (a) Topology diagrams for the GT1 (teal) and GT2 (gold) domains, respectively, drawn using TOPDRAW [1]; (b) Superposition of structural homologs to the GT1 domain of GLT25D1/COLGALT1 (shown as thick black ribbon), as identified using DALI with Z-score higher than 15. The table inset indicates the PDB ID of each entry, as well as the associated DALI Z-scores and RMSD values. (c) The closest structural homolog of GLT25D1/COLGALT1 GT1 domain is the AC domain of human LH3/PLOD3. For reference, the image shown on the right hightlights the position of the AC domain within the complete LH3/PLOD3 polypeptide (PDB ID 6FXR). The superposition shows the excellent match between the two molecular architectures (shown in teal for GLT25D1/COLGALT1 and orange for LH3/PLOD3, respectively) and the key differences (shown with a black arrow) in the region proximate to the cofactor and donor substrate binding site identified in GLT25D1/COLGALT1. (d) Superposition of structural homologs to the GT2 domain of GLT25D1/COLGALT1, as identified using DALI with Z-score higher than 10. The table inset indicates the PDB ID of each entry, as well as the associated DALI Z-scores and RMSD values.

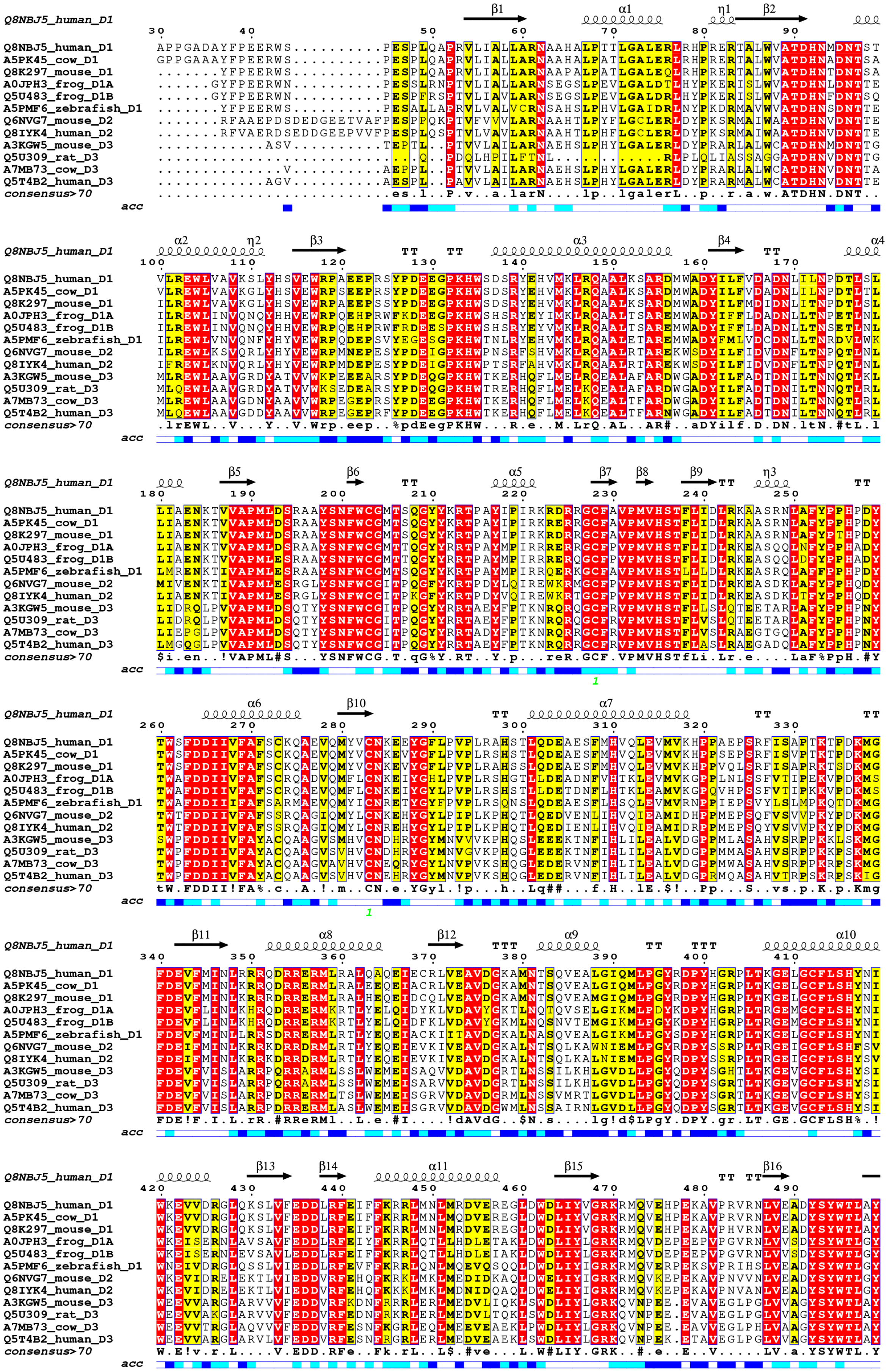
Q8NBJ5_human_D1
A5PK45_cow_D1
Q8K297_mouse_D1
A0JPH3_frog_D1A
Q5U483_frog_D1B
A5PMF6_zebrafish_D1
Q6NVG7_mouse_D2
Q8IYK4_human_D2
A3KGW5_mouse_D3
Q5U309_rat_D3
A7MB73_cow_D3
Q5T4B2_human_D3
consensus>70
acc

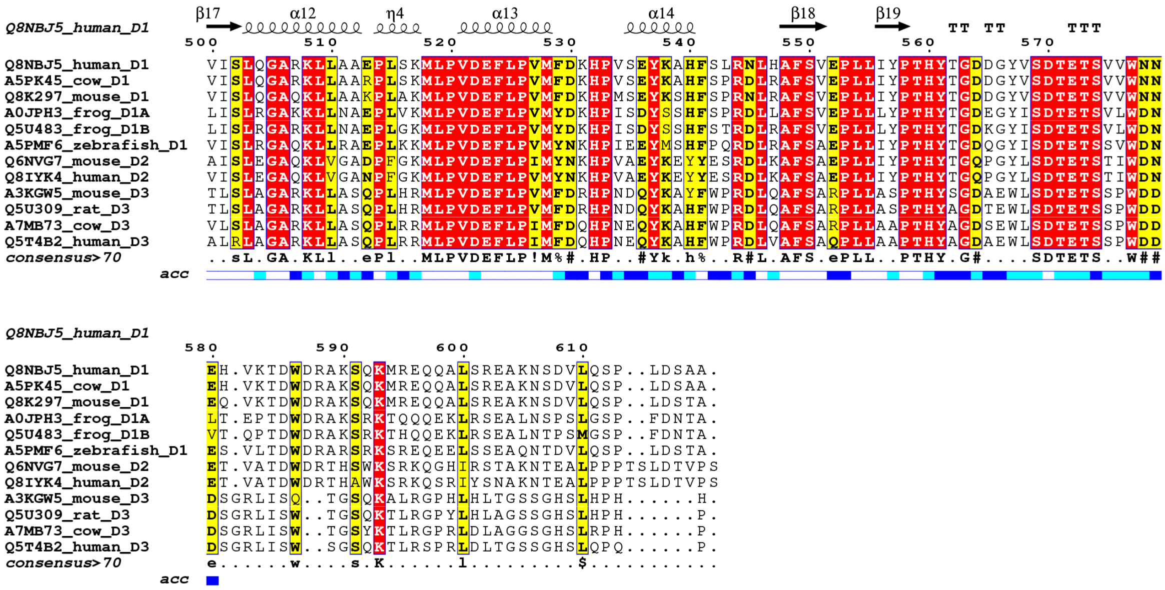


**Supplementary Figure 4. Multiple sequence alignment of GLT25D/COLGALT sequences from different animal species.** The alignment has been generated using EBI MUSCLE[2] and drawn using ESRIPT[3]. The "*consensus*" line computes the sequence consensus according to the result of the multiple sequence alignment (global score 0.7, differential score 0.5). The "*acc*" line indicates the solvent accessibility of each residue based on evaluation of the experimental structure of human GLT25D1/COLGALT1 (white, non-accessible residues; blue, accessible residues).

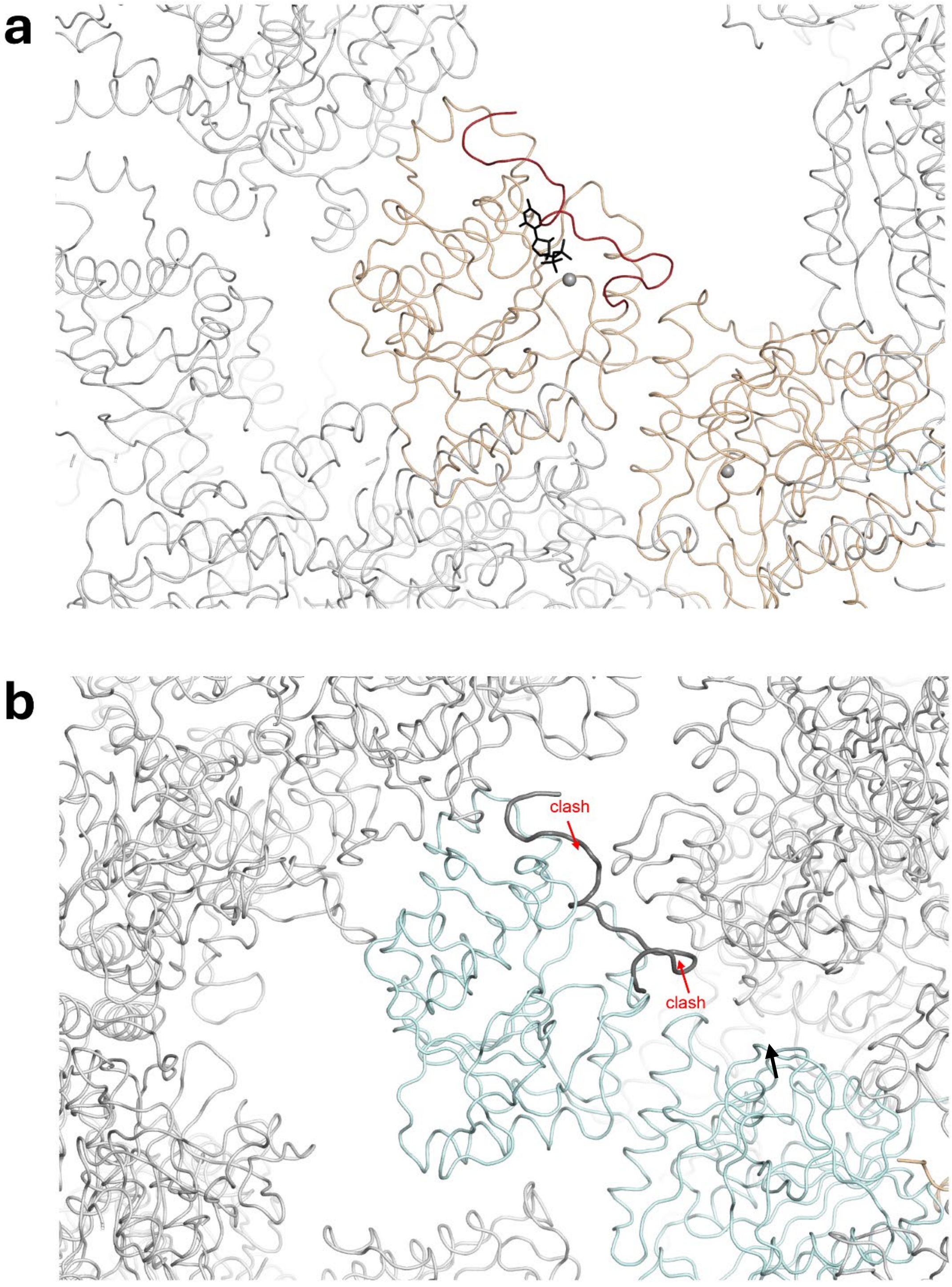


**Supplementary Figure 5. Overview of the crystal packing observed in the crystal structures of GLT25D1/COLGALT1.** The representations highlight the different contacts surrounding the GT2 domains in the two copies of the enzyme found in the asymmetric unit of the enzyme structure obtained with excess $Mn^{2+}$ and UDP-$\alpha$-Gal. The conformation adopted by the 560-570 loop in the substrate-bound structure, compatible with the packing of monomer A shown is panel (a), is not compatible with the crystal packing of the second monomer (b). In B, the loop from superposed monomer A is shown as thick grey ribbon to highlight the packing clashes.

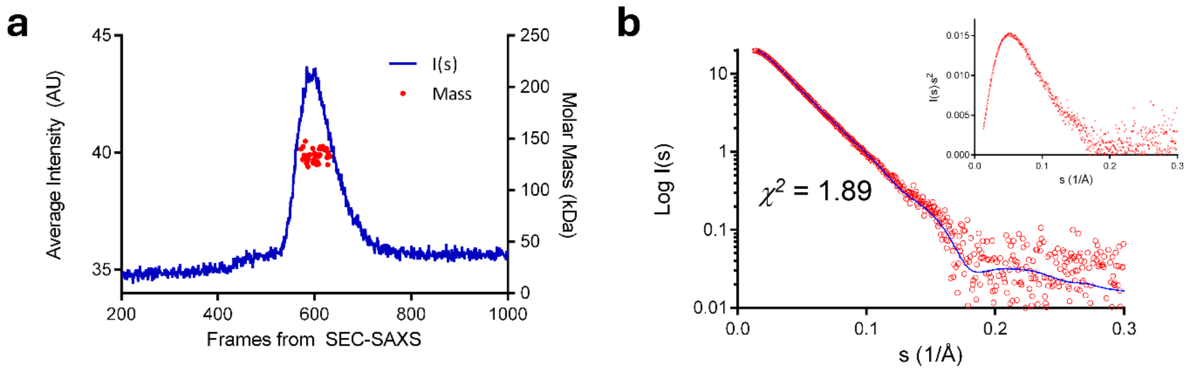


**Supplementary Figure 6. Results of SEC-SAXS analysis of GLT25D1/COLGALT1 in solution.** (a) SEC-SAXS chromatogram and molecular weight analysis of peak fractions derived from CHROMIXS analysis[4]. (b) SAXS data obtained from the analysis of the peak frames of the SAXS chromatogram shown in (a) (red circles, with Kratky transformation in the inset), overlaid to the computed SAXS profile (solid blue line) from the *CORAL* model, shown in Figure 3D, resulting in fit agreement with $\chi^2$ of 1.89.

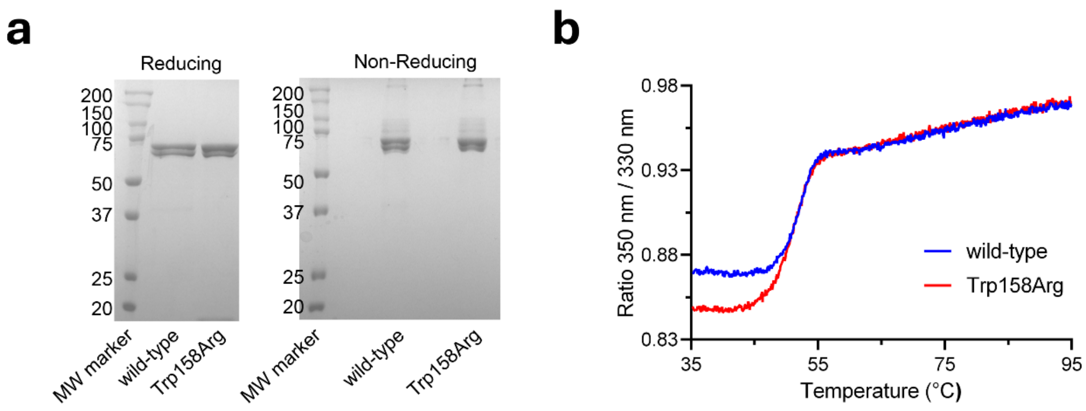


**Supplementary Figure 7. Quality control for the GLT25D1/COLGALT1 dimer interface mutant Trp158Arg.** The mutant generated underwent testing for its correct folding using reducing and non-reducing SDS-PAGE (a) and DSF analysis (b).

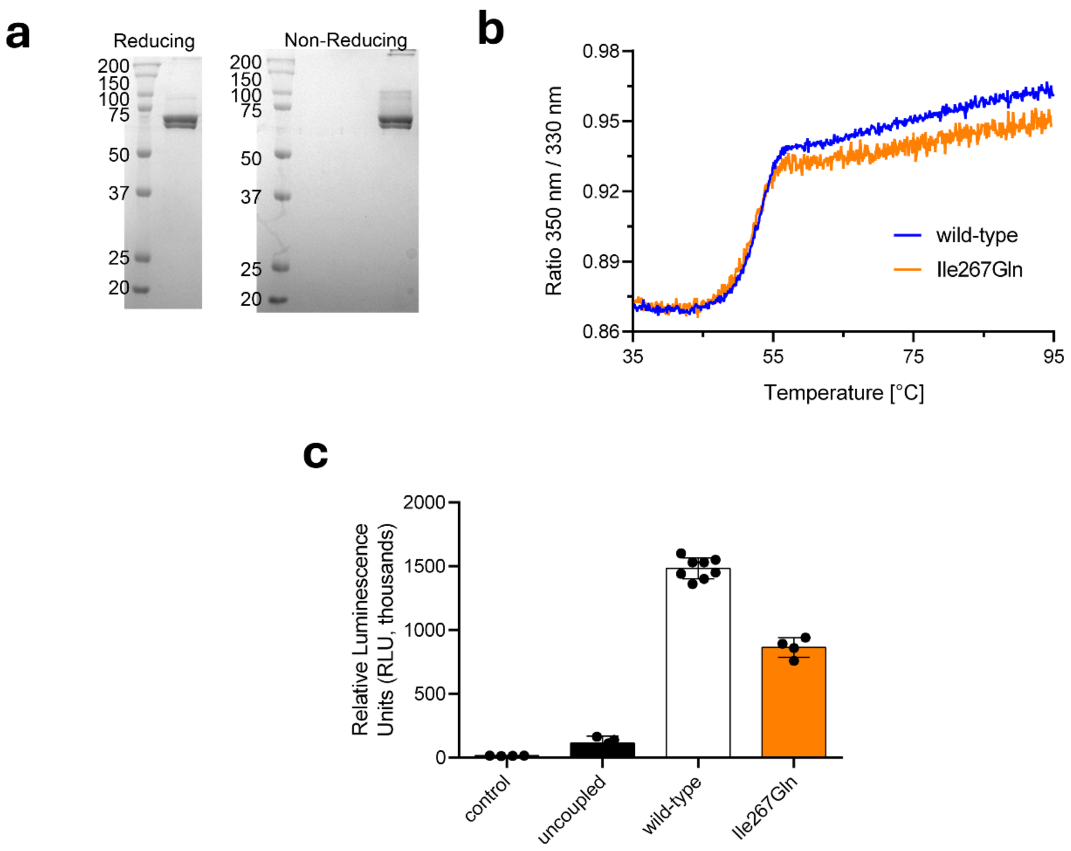


**Supplementary Figure 8. Characterization of GLT25D1/COLGALT1 Ile267Gln mutant.** All purification attempts focusing on GLT25D1/COLGALT1 GT1 mutants involved in UDP-Gal binding resulted in no expression or improperly folded enzyme preparations. Conversely, the control mutant Ile267Gln, proximate to UDP-Gal binding in the GT1 domain but not directly involved in cofactor binding, could be purified, as confirmed by SDS-PAGE (a) and DSF analysis (b). (c) The enzymatic activity of control mutant Ile267Gln has been compared with wild-type enzyme using luminescence-based activity assays.

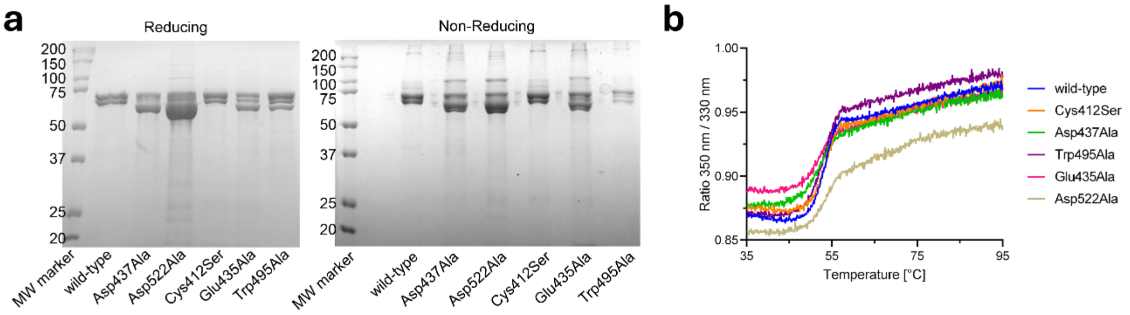


**Supplementary Figure 9. Quality control for the GLT25D1/COLGALT1 GT2 catalytic site mutants.** The mutants generated underwent testing for its correct folding using reducing and non-reducing SDS-PAGE (a) and DSF analysis (b).
